# A systems based framework to computationally predict putative transcription factors and signaling pathways regulating glycan biosynthesis

**DOI:** 10.1101/2020.08.19.257956

**Authors:** Theodore Groth, Sriram Neelamegham

**Affiliations:** Chemical and Biological Engineering, State University of New York, Buffalo, NY 14260, USA; Biomedical Engineering, State University of New York, Buffalo, NY 14260, USA; Medicine University at Buffalo, State University of New York, Buffalo, NY 14260, USA

**Keywords:** Glycoinformatics, transcription factor, glycosylation, ChIP-Seq, TCGA

## Abstract

Glycosylation is a common post-translational modification, and glycan biosynthesis is regulated by a set of ‘glycogenes’. The role of transcription factors (TFs) in regulating the glycogenes and related glycosylation pathways is largely unknown. This manuscript presents a multi-omics data-mining framework to computationally predict putative, tissue-specific TF regulators of glycosylation. It combines existing ChIP-Seq (Chromatin Immunoprecipitation Sequencing) and RNA-Seq data to suggest 22,519 potentially significant TF-glycogene relationships. This includes interactions involving 524 unique TFs and 341 glycogenes that span 29 TCGA (The Cancer Genome Atlas) cancer types. Here, TF-glycogene interactions appeared in clusters or ‘communities’, suggesting that changes in single TF expression during both health and disease may affect multiple carbohydrate structures. Upon applying the Fisher’s exact test along with glycogene pathway classification, we identify TFs that may specifically regulate the biosynthesis of individual glycan types. Integration with knowledge from the Reactome database provided an avenue to relate cell-signaling pathways to TFs and cellular glycosylation state. Whereas analysis results are presented for all 29 cancer types, specific focus is placed on human luminal and basal breast cancer disease progression. Overall, the computational predictions in this manuscript present a starting point for systems-wide validation of TF-glycogene relationships.

## Introduction

The glycan signatures of cells and tissue is controlled by the expression pattern of 300-350 glycosylating genes that are together termed ‘glycogenes’ [1,2]. These glycogenes include the glycosyltransferases, glycosidases, sulfotransferases, transporters etc. The expression of these glycogenes is in turn driven by the action of a class of proteins called transcription factors (TFs). These TFs regulate gene expression by binding proximal to the promoter regions of genes, facilitating the binding of RNA polymerases. They may homotropically or heterotropically associate with additional TFs in order to directly or indirectly control messenger RNA (mRNA) expression. Among the TFs, some ‘pioneer factors’ can pervasively regulate gene regulatory circuits, and access chromatin despite it being in a condensed state [3]. These TFs act as ‘master regulators’, promoting the expression of several genes across many signaling pathways, such as differentiation, apoptosis and cell proliferation. The precise targets of the TFs is controlled by their tissue-specific expression, DNA binding domains and nucleosome interaction sequences [3]. Additional factors regulating transcriptional activity include: i. cofactors and small molecules that enable TF-DNA recognition and RNA polymerase recruitment [3]; ii. chomatin modifications, such as acetylation, methylation and phosphorylation, which alter TF access; and iii. methylation of CpG islands in promoter regions which inhibit gene expression [4,5].

There are currently several isolated studies of TF-glycogene interactions, but to our best knowledge a systematic “systems level analysis” is absent. Many of these previous studies are based on discrete glycogene promoter region analysis and reporter assays. These studies have established some notable TF-glycogene relationships, though they are limited to distinct cell types. Examples include the regulation of MGAT5 by ETS2 in NIH3T3 fibroblasts [6], control of the α2-6 sialyltransferases ST6Gal-I/II by Hypoxic Nuclear Factor 1-α (HNF1-α) in HepG2 cells [7], c-JUN-B3GNT8 regulatory relationships in gastric carcinoma cell lines [8], and SP1-B4GALT1 relations in lung cancer A549 cells [9]. A recent study also used computational predictions and wet-lab experiments to determine that ZNF263 is a potential heparin sulfate master regulator [10]. This TF regulates two sulfotransferases, HS3ST1 and HS3ST3A1. The above approaches have limitations in that: i. they do not consider the cellular epigenetic state that could impact TF binding; and ii. proximal regulators are studied but enhancers present several kilobases away from the <u>T</u>ranscription <u>S</u>tate <u>S</u>ite (TSS) away are neglected.

In the current manuscript, we propose that more global and higher-throughput TF-glycogene relationships under biologically relevant conditions may be discovered using multiomics data-mining. To this end, we sought to utilize multi-omics experimental datasets and curated pathway databases (DB) to relate cell-specific signaling processes to TFs, TFs to glycogenes, and glycogenes to glycosylation pathways (**Fig. 1A**). These connections were made using data available from Cistrome Cancer DB [11], Reactome DB [12] and by the manual curation of various human glycogenes into pathways at GlycoEnzDB (www.virtualglycome.org/GlycoEnzDB, **Fig. 1B**). Here, the Cistrome Cancer DB uses TF-gene binding data from previously published ChIP-Seq studies for various cell systems and cancer tissue RNA-seq data from The Cancer Genome Atlas (TCGA) [13]☐. It curates putative TF-gene relationships for 29 TCGA cancer types provided they satisfy three inclusive criteria: i. TFs should be expressed at high levels in a given tissue; ii. Changes in TF gene expression should correlate with RNA changes in target genes; iii. Chromatin Immunoprecipitation sequencing (ChIP-Seq) data must support the TF-gene binding proximal to the transcription start site (TSS). Next, knowledge curated in the Reactome DB [12]☐ is used to establish links between TFs and signaling pathways. In the final step, manually curated glycogene classifications were utilized to determine TFs that disproportionately regulate individual glycosylation pathways. The findings from this study Clusters in the resulting interactomes were related to pathway maps and signaling processes, thus developing TF-community signaling pathway relationships.

**Figure 1.**
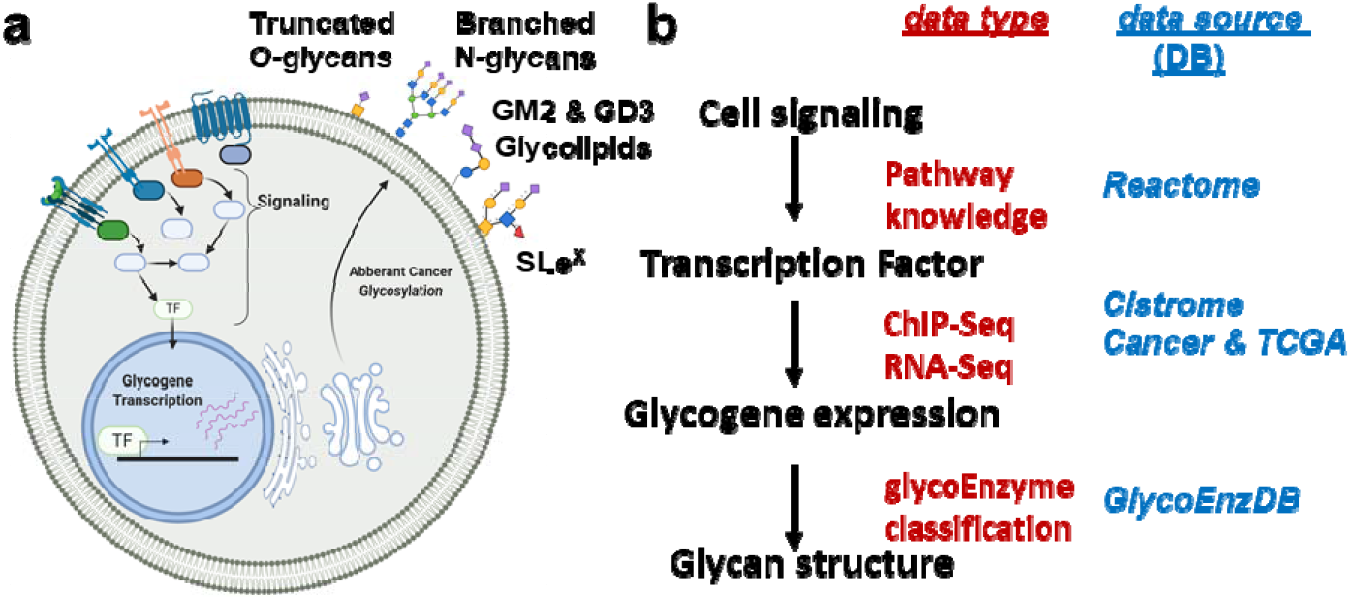
A systems glycobiology framework to link multi-OMICs data: **a.** Cell signaling proceeds to trigger transcription factor (TF) activity. The binding of TFs to sites proximal to the transcriptional start site triggers glycogene expression. A complex set of reaction pathways then results in the synthesis of various carbohydrate types, many of which are either secreted or expressed on the cell surface. **b.** Data available at various resources can establish the link between cell signaling and glycan biosynthesis. The Reactome DB contains vast cell signaling knowledge. Chip-Seq and RNA-Seq data available at the Cistrome Cancer DB describe the link between the TFs and glycogenes. Pathway curation at the GlycoEnzDB establishes the link between glycogenes and glycan structures. Cell illustration created using BioRender.com.

## Results

### TF-Glycogene interaction map and relation to cell signaling pathways

The manuscript follows a workflow shown in **Figure 2**. It infers TF-glycosylation pathway relationships, as well as TF-glycogene communities by using publicly available ChIP-seq data and RNA-Seq results. These data were obtained from the Cistrome Cancer DB [14], which curated TF-gene relationships by integrating ChIP-seq data from Cistrome DB and RNA-seq data from The Cancer Genome Atlas (TCGA). The Cistrome Cancer DB uses three filtering criteria to determine putative TF-gene relationships: 1. The TF should be active in a cancer type, i.e. the RPKM (<u>R</u>eads <u>P</u>er <u>K</u>ilobase <u>M</u>illion) value in a cancer type must be greater than the TF’s median RPKM expression across all 29 different cancer types. 2. The RNA expression of TF and target gene should be correlated. To determine this, Cistrome first compares the selected TF-gene correlation with a null distribution computed by randomly selecting 1 million TF-gene pairs. Linear regression and statistical analysis are then performed on the top 5-percentile hits (positive and negative coefficients) to establish TF-gene correlations. This analysis accounts for target gene copy number, tumor purity, and promoter methylation extent. 3. TF-gene relationships must be supported by Chromatin Immunoprecipitation sequencing (ChIP-Seq) evidence. Here, a non-linear weighted sum called “regulatory potential (RP)” quantifies the strength of TF-gene interactions based on the proximity of TF binding site to the gene transcriptional start site, and also the number of TF-gene binding interactions based experimentally detected ChIP-peaks [15,16].

**Figure 2.**
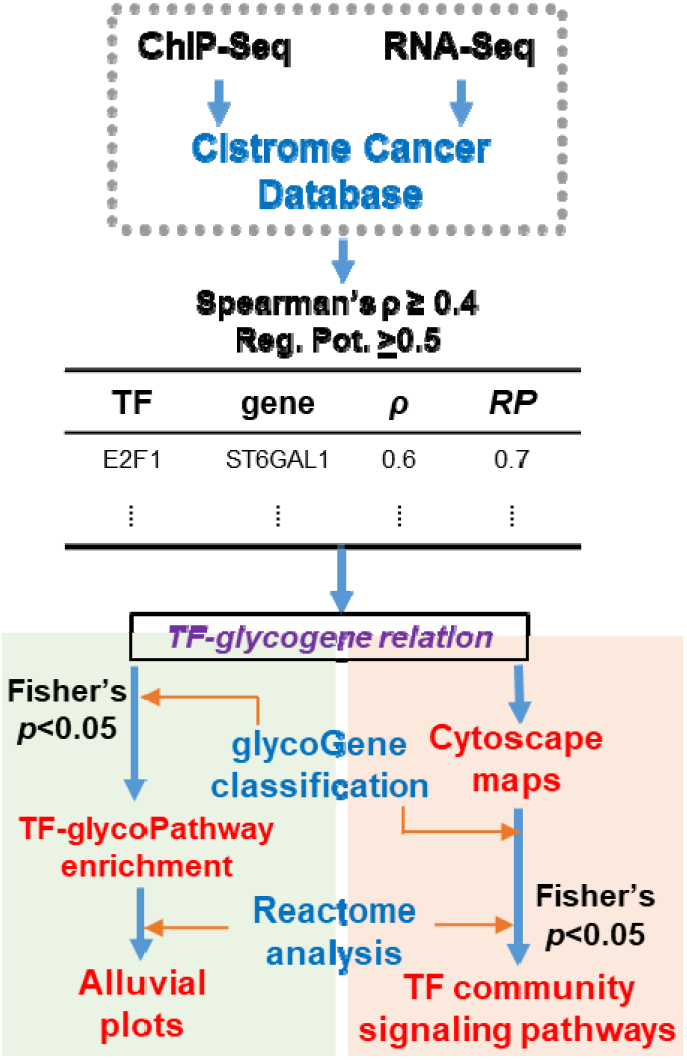
Analysis workflow: ChiP-seq provides evidence of TF binding to promoter regions with regulatory potential (0≤*RR*≤1) quantifying the likelihood that this is functionally important. RNA-Seq quantifies Spearman’s correlation (ρ) between TF and gene expression. Filtering these data establishes potential TF-glycogene interactions in specific cancer types. TFs disproportionately regulating specific glycosylation pathways were identified using the above TF-glycogene relationships, and biochemical knowledge available at GlycoEnzDB (green region). ReactomeDB analysis helped establish cell signaling-TF-glycosylation pathway connectivity that are visualized using Alluvial plots. Independently, cytoscape maps enabled visualization of TF-glycogene relationships in different cancers (orange region).

In the current manuscript, we passed the TF-gene relationships established in Cistrome Cancer DB, to identify TFs potentially interacting with 341 glycogenes (**Supplemental Table S1**). The two metrics for this select were RP≥0.5 and TF-glycogene expression correlation coefficient ρ≥0.4. Such analysis was performed for 29 cancer types listed in **Supplemental Table S2**. Based on our selected thresholding, the analysis revealed 22,519 potential TF-glycogene interactions. The above data were used for two types of analysis described below.

First, the Fisher’s exact test was used to infer TF-glycogene interactions that may regulate individual glycosylation pathways. This analysis was based on pathway classifications from GlycoEnzDB (**Supplemental Table S3**) that grouped 208 glycogenes into 20 glycosylation pathways/groups. TFs having disproportionately larger number of relationships with individual glycosylation pathways were determined, with respect to all TF-glycogene relationships. Reactome DB was then used to associate these TFs to potential signaling pathways. This method, thus, resulted in a relationship between cell-signaling, TF activity regulation and glycan structure changes (**Supplemental Table S4, S5**). The data are presented as Alluvial plots for the 29 cancer types (**Supplemental Fig. S1**). Here, the TFs were linked to glycosylation pathways by colored bands if they were found to regulate a disproportionately high fraction of glycogenes belonging to that pathway. Likewise, biological pathways were linked with TFs if that TF was found to be enriched in the biological pathway. Reading these alluvial plots left to right, one can deduce which biological pathways may be potentially involved in regulating TFs, and how these TFs could regulate glycosylation.

Second, we visualized TF-glycogene interactions using Cytoscape maps for each of the cancers individually (**Supplemental File S2**). This analysis revealed clusters of TFs regulating common sets of glycogenes. We sought to understand which glycosylation pathways were overrepresented in each of these cluster using the Fisher’s exact test. This analysis revealed 406 glycopathway enrichments in the TF-glycogene communities across the 29 cancer types (**Supplemental Table S6**). Next, we determined, using the Reactome DB overrepresentation API, if the TFs identified in these clusters could be related to specific cell signaling pathways. Here we noted 901 pathway enrichments across the different cancer types (**Supplemental Table S7**). Common TFs that we observed across all TF-glycogene communities include the TCF and LEF families, FOXO and FOXP, RUNX family, and IRF family TFs, which were found to regulate diverse glycosylation pathways, such as sialylation pathways, complex N-linked glycan synthesis, chondroitin and dermatan sulfate synthesis.

Overall, the above analysis revealed the existence of communities of TF-glycogene relationships, that could be linked to both cell signaling processes and specific glycosylation pathways. Future, quantitative computational modeling and mass spectrometry studies may establish the biological significance of these predictions.

### TF-pathway relationships in breast cancer

As a usage example of the above framework, we provide more detailed investigation using breast cancer as an example. This disorder appears in 5 unique molecular subtypes based on the PAM50 classification [17]. These include: i. normal-like, ii-iii. luminal A and luminal B which overexpress estrogen receptor ESR1, iv. Her2+ tumors that overexpress the epidermal growth factor receptor (ERBB), and v. basal (triple negative) that express neither ESR1 nor ERBB. Each of these subtypes has unique signaling mechanisms that may contribute to different glycan signatures.

In our analysis, TF-glycogene relationships for breast cancer derived by filtering Cistrome Cancer were enriched for the glycosylation pathways. **Fig. 3** summarizes these cancer related TF-glycosylation pathway relationships for luminal (type A and B together) and basal breast cancers. The analysis suggests that TF transformations accompanying cancer progression may impact all four major classes of glycans: O-and N-glycans found on glycoproteins, glycosaminoglycans and glycolipids. Thus, multiple glycan changes may accompany oncological transformation.

**Figure 3.**
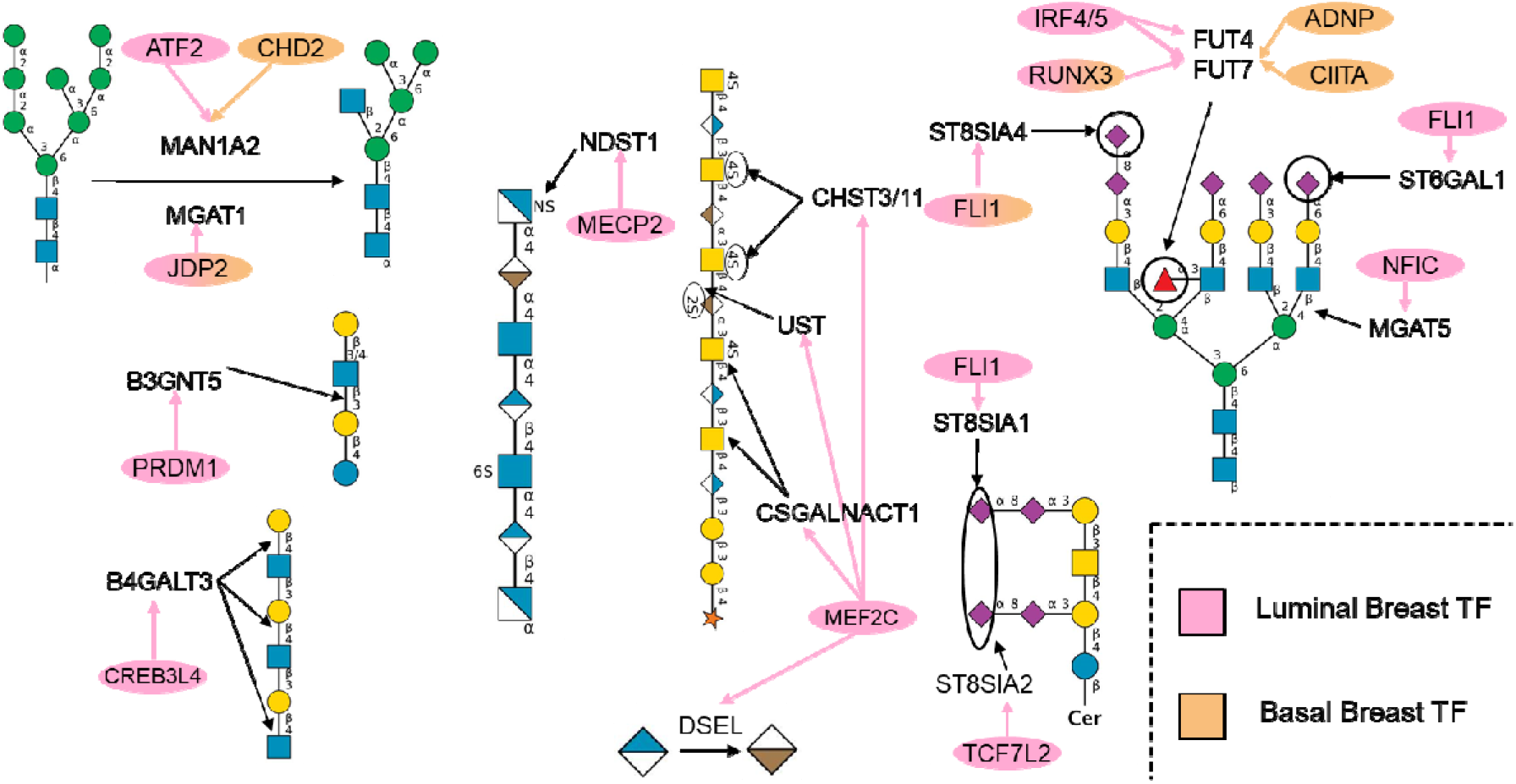
Summary of TFs enriched to glycosylation pathways for luminal and basal breast cancer: The TFs found to be enriched to glycosylation pathways and the glycogenes they regulate are shown in pink for luminal and orange for basal breast cancer. Note that some of the TFs shown above do not appear in the alluvial plots in figures 4 and 5 because they were not enriched to a signaling pathway in Reactome. The glycans synthesized by the enriched glycogenes are shown in SNFG format [18]. All figures were generated using DrawGlycan-SNFG [19].

### TF-glycogene communities in luminal and basal breast cancer (cytoscape plots)

Cytoscape plots were generated for luminal breast cancer (**Fig. 4a**). Here, using the bipartite graph community detection methods [20], we inferred three large communities of TF-glycogene interactions. The largest community detected in this analysis had TFs enriched for RUNX3 signaling, IL-21 signaling, MECP2, and PTEN regulation. Over-representation glycosylation pathway analysis performed on the TFs in this community suggests that these TFs may regulat**e** pathways related to sialylation, hyaluronan synthesis, and chondroitin and dermatan sulfate elongation. Here, STAT1, 4, and 5 proteins were enriched in the IL-21 signaling pathway. Luminal breast cancers are known to express STAT1, 3 and STATs 2 and 4 are known to be expressed in luminal breast cancer cell lines. STAT5 is known to be constitutively active in luminal breast cancer and confers anti-apoptotic characteristics to cells [21]. The other two communities detected consisted primarily of chromatin-modifying enzymes. Complex N-linked glycan synthesis and the dolichol pathway were significantly enriched in the second community.

**Figure 4.**
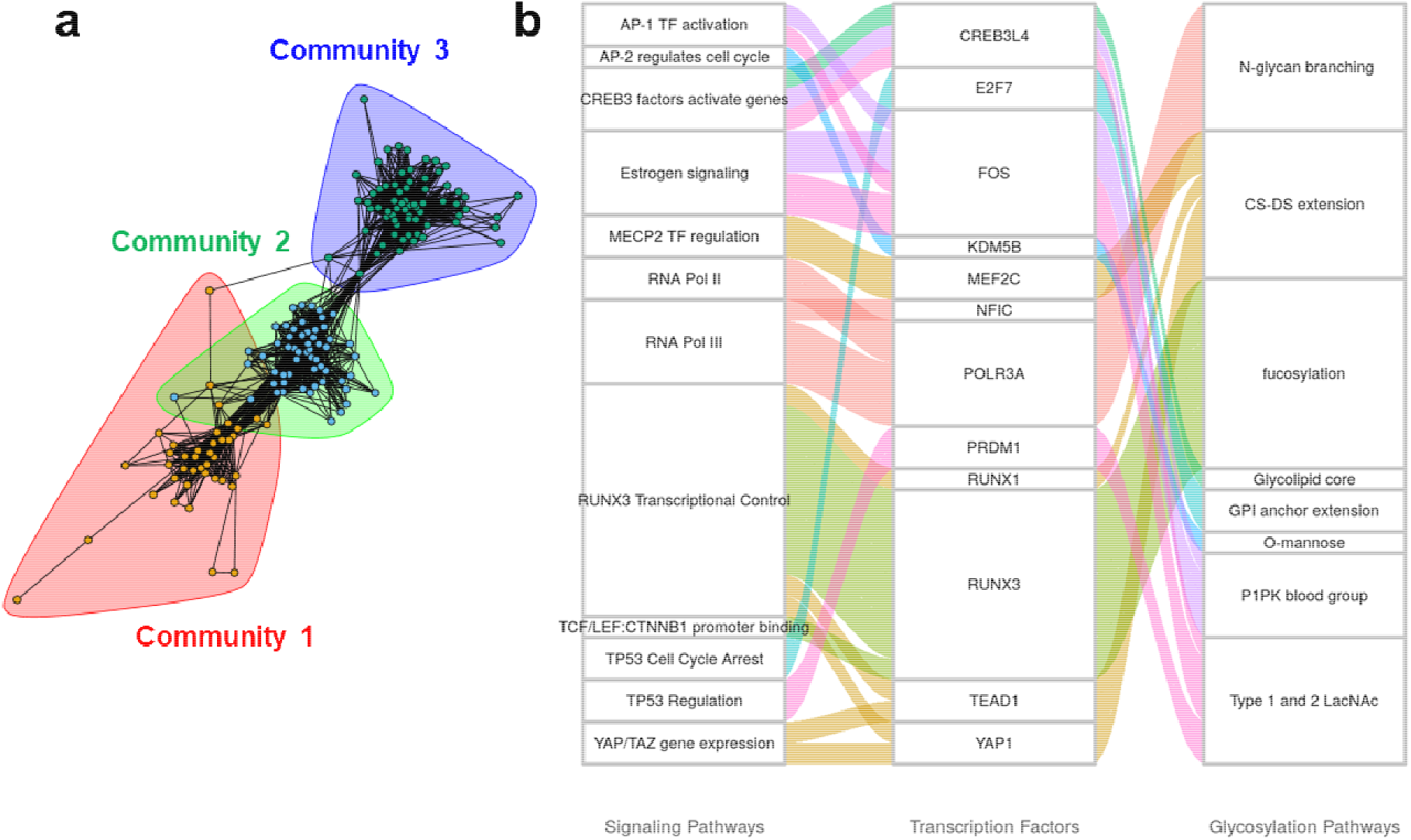
Luminal breast cancer signaling pathway enrichment and glycogene connections: **a.** *TF-to-glycogene communities in luminal breast cancer:* Three large TF-to-glycogene communities were discovered in the luminal breast subnetwork. Community 1 was enriched for pathways involving RUNX3, RUNX1, IL-21, and PTEN. Communities 2 and 3 consist primarily of chromatin modifying enzymes. **b.** *Signaling pathway enrichment analysis for luminal breast cancer*: Connections between signaling pathways and transcription factors found to be statistically significant for luminal breast cancer. Some pathways enriched to TFs were condensed to conserve space. More TF-to-glycogene relationships exist in luminal breast cancer and these can be viewed in the cytoscape figures (**Supplemental Figure S2**).

In the third community, O-linked mannose and LacdiNAc synthesis were disproportionately regulated. Overall, the pathway maps suggest that chromatin remodeling enzymes could potentially play roles in regulating glycan synthesis in luminal breast cancer.

Like luminal, basal breast cancer TF-glycogene relationships were also clustered into three communities. Here, the first community was enriched for chromatin modifying enzymes, with complex N-linked glycan synthesis being the primary glycosylation pathway being affected (**Fig. 5a**). The second community was enriched for interferon *α/β/γ* signaling pathways, with interferon regulatory factor (IRF) TFs being enriched. In this regard, the TFs IRF-1 and IRF-5 have been shown to act as tumor suppressors in breast cancer [22,23]. Their loss-of-function in breast cancer could potentially downregulate O-linked fucosylation. The third community did not exhibit any specific TF pathway enrichments.

**Figure 5.**
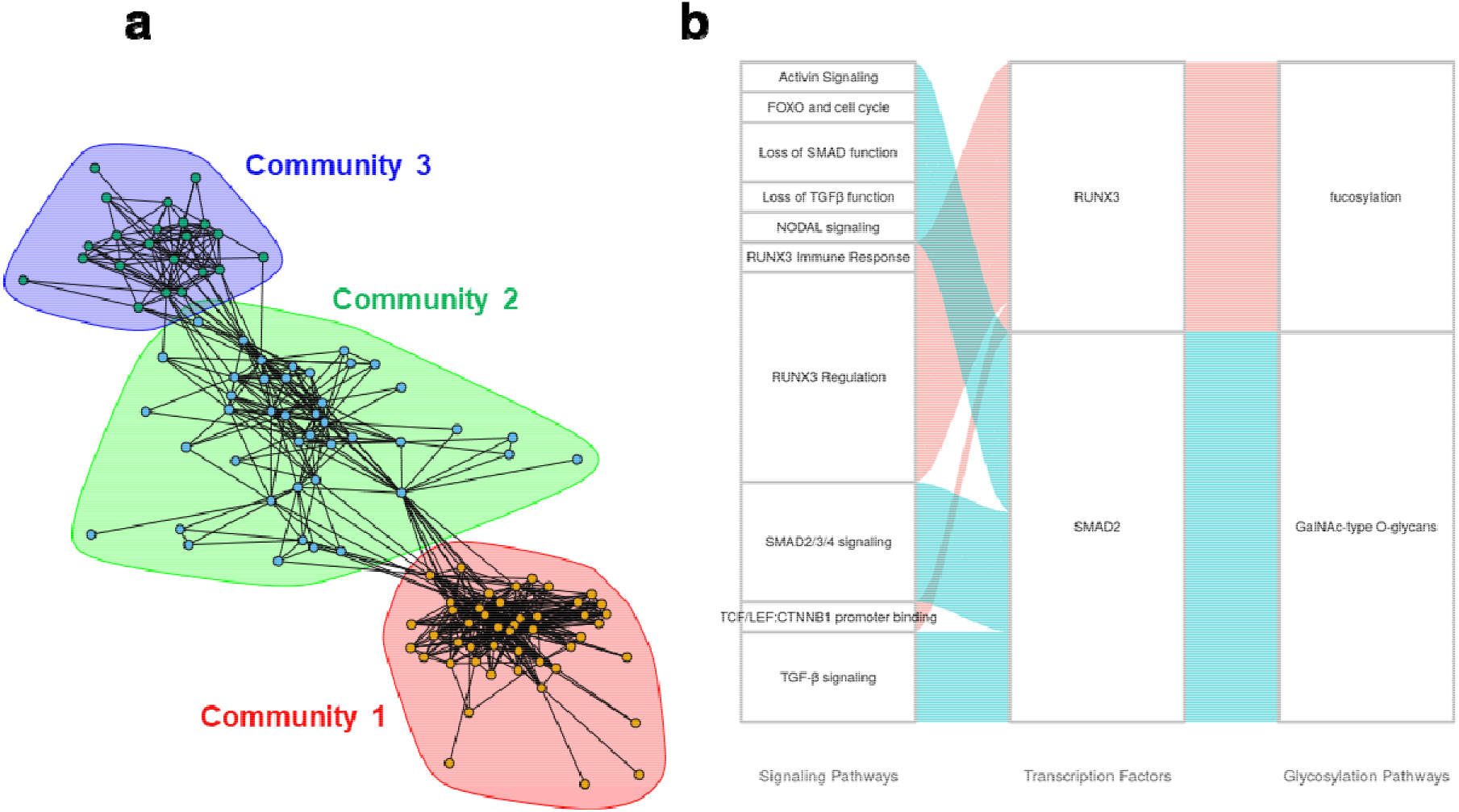
Basal breast cancer signaling pathway enrichments and glycogene connections: **a.** *TF-to-glycogene communities in basal breast cancer:* Three large TF-to-glycogene communities were discovered in the basal breast subnetwork. Community 1 has TFs enriched to chromatin modifying enzymes, and community 2 has TFs enriched to interferon α/β/γ signaling. Community 3 did not have any signaling pathways enriched. **b.** *Signaling pathway enrichment analysis for basal breast cancer*: Connections between signaling pathways and TFs found to be statistically significant for basal breast cancer. TFs displayed have been enriched to the displayed glycosylation pathways using the Fisher’s exact test.

### Linking cell signaling to TF and glycogenes for luminal breast cancer (alluvial plot)

The links between biological signaling pathways, TFs, and glycosylation pathways are shown in alluvial plots for luminal (**Fig. 4b**) and basal breast cancers (**Fig. 5b**), with additional plots provided for additional cancer types in Supplemental Material. In the case of luminal breast cancer:

#### CREB3L4 and PRDM1 disproportionately affect the Type I & II LacNAc pathway in luminal breast cancer

Our analysis suggests that CREB3L4 (enrichment p-value = 0.036) and PRDM1 (enrichment p-value = 0.039) may regulate the Type 1 and 2 LacNAc pathways. CREB3L4 is known to primarily be expressed in the prostate and some breast cancer cell lines, and has been linked to diverse roles involving chromatin organization in spermiogenesis, adipocyte regulation, and dysregulation in prostate cancer[24,25]. It has been found to be upregulated in breast cancer with respect to normal. PRDM1, also known as Blimp-1, is a transcriptional repressor, and its upregulation in cancer is known to dysregulate other proteins [26].The increase Poly LacNAc structures have been shown to play roles in cancer metastasis [27]. CREB3L4 was found to regulate B4GALT3 glycogene (ρ=0.56, RP=0.94) which adds galactose in a β1-4 linkage, and PRDM1 was found to regulate B3GNT5 which is critical for lacto- /neolactoseries glycolipids (ρ=0.60, R.P.=0.84).

#### MEF2C disproportionately regulates Glycosaminoglycan synthesis pathways

MEF2C was found to regulate several genes in the chondroitin and dermatan sulfate synthesis pathways (p=0.008). This TF plays roles in development, particularly with the development of neurons and hematopoetic cell differentiation towards myeloid lineages. It is known that MEF2C is directly impacted by TGF-β signaling, thus increasing the metastatic potential of cancer [28]. MEF2C was found to be inhibited by MECP2 based on Reactome pathway enrichment. Since the glycosaminoglycan elongation pathways positively correlate to MEFC2 expression, and MEFC2 is amplified in cancer, it is possible that MECP2 may not sufficiently expressed to repress MEFC2 in call cancer cells. MEF2C was found to regulate CSGALNACT1 (ρ=0.66, RP=0.71), CHST3 (ρ=0.50, RP=0.74), CHST11 (ρ=0.47, RP=0.84), DSEL (ρ=0.40, RP=0.81), and UST (ρ=0.42, RP=0.95). Here, CSGALNACT1 is responsible for the addition of GalNAc to glucuronic acid to increase chondroitin polymer length, CHST3, CHST11, and UST are involved in the sulfation of GalNAc and iduronic acid, and DSEL is the epimerase which converts glucuronic acid to iduronic acid in CS/DS chains.

#### MECP2 disproportionately regulated heparan sulfate chain elongation

The MECP2 (enrichment p-value=0.037) was found to positively regulate heparan sulfate elongation. MECP2 regulates gene expression by binding to methylated promoters, and then by recruiting chromatin remodeling proteins to condense DNA and repress gene expression [29,30]. MECP2 was found to regulate sulfotransferase NDST1 (ρ=0.41, RP=0.67).

### Linking cell signaling to TF and glycogenes for basal breast cancer (alluvial plot)

Fewer transcription factors were found to be enriched to signaling pathways in basal breast cancer compared to luminal cancer (**Fig. 4b**). Despite this, there are many other TF-glycosylation pathway enrichments for basal breast cancer available for analysis in Supplementary Material. The roles of the enriched TFs and their relation to glycogenes and cancer is elaborated below.

#### RUNX3 and Fucosylation

The terminal fucosyltransferase FUT7 (ρ=0.49, RP=0.89) was found to be positively regulated by the RUNX3 TF (enrichment p-value = 0.033). The RUNX family of transcription factors (including RUNX1-3), are involved in several developmental processes, including hematopoiesis, immune cell activation, and skeletal development. It was discovered that RUNX3 acts as a tumor suppressor gene in breast cancer. Upon cancer development, the RUNX3 promoter is hypermethylated, leading to reduced TF activity and loss of tumor suppression activity [31]. Our data suggest that this may be associated with a reduction of FUT7 activity thus impacting the expression of the sialyl Lewis-X antigens in basal tumors. Sialyl Lewis-X is considered to be an important regulator of cancer metastasis as it binds the selectins on various vascular and blood cell types.

#### Regulation of GalNAc-type O-linked glycans by SMAD2

SMAD2 was found to significantly affect core 1 & 2 O-linked glycan structures (enrichment p-value=0.035). SMAD proteins are activated by TGF-β signaling and bind to DNA to act as cofactors to recruit TFs. SMAD2 has been shown to act as a tumor metastasis suppressor in cell lines [32,33]. This TF was found to regulate GALNT1 (ρ=0.54, RP=1.00), which adds GalNAc to serine or threonine residues to being core 1 and 2 O-linked glycan synthesis. Thus, SMAD2 may play a key role in regulating Tn-antigen expression in proteins like MUC-1 that are associated with breast cancer progression.

## Discussion

In the current analysis, we sought to identify strategies to enhance systems glycobiology knowledge by leveraging existing, publicly-available, high-throughput ChIP-Seq and RNA-Seq data. This analysis suggests putative TFs regulating glycogenes and glycosylation processes across 29 different cancer types. Approximately three glycogenes were regulated by a given TF based on our filtering criteria, with this number ranging from 1-10. These findings are tissue-specific, as TF and glycogene expression vary widely among the different cell types. The analysis also suggests putative TF-glycogene interactions that disproportionately impact specific glycosylation pathways. Knowing which TF regulates which glycogene and pathway in a context-dependent manner can provide insight as to how signaling pathways contribute to altered glycan structures in diseases such as diabetes and cancer. Thus, this work represents a rich starting point for wet lab validation and glycoinformatics database construction.

Visualizing TF-glycogene interaction networks revealed communities of glycogenes in each cancer type. The presence of chromatin-modifying enzymes in large regulatory communities in both luminal and basal breast cancer suggests a role of epigenetics in glycogene regulation. To date, a systems-level investigation evaluating the epigenetic states of cell systems on the resulting glycome has not been performed. Our results suggest that complex N-linked branching and glycosylation may be sensitive to these processes. The signaling pathways enriched in the largest community in luminal breast cancer were reflected in our pathway enrichment findings. RUNX3, interleukin signaling, and the involvement of MECP2 regulation were all found to disproportionately regulate sialic acid and GAG synthesis pathways.

Several of the TFs enriched to glycosylation pathways were either regulated by or involved in TGF-β signaling and Wnt β-catenin signaling. These TFs primarily affected glycosaminoglycan synthesis pathways, sialylation and Type-2 LacNAc synthesis. Cell cycle and metabolic regulatory TFs were shown to regulate some glycogenes involved in the dolichol pathway. The crosstalk between cell cycle and glycosylation is not well explored, and may potentially be important for understanding N-linked glycosylation flux in cancer. Some TFs were found to interact with methyl CpG binding TFs when regulating glycosaminoglycan proteins, implicating methylation as a possible modulator of glycosylation in cancer.

The framework described in this manuscript represents a starting point for experimentally investigating TFs regulating glycosylation, however much more is needed to validate and establish these relationships. First, selected values of RP and ρ used to filter TF-glycogene relationships from the Cistrome Cancer database were selected to initiate a query into the available ChIP-seq data. Further studies are needed in order to determine how the selected thresholds affect the discovered relationships. Second, the glycogenes in individual pathways in this manuscript were classified using current knowledge of glycobiology. Different classification methods may result in slightly different TF-glycogene relationships [34]. Third, while Cistrome Cancer DB systematically filters TF-gene relationships based on ChIP-seq and RNA-seq evidence, the database has some biases. In one aspect, only TFs that were considered to be sufficiently expressed were considered in this analysis. Lower expressed TFs that may also be functional, are excluded from the analysis. Additionally, while RNA-Seq relationships in Cistrome Cancer are selected based on specific tissue type, supporting ChIP-seq evidence is not cell type specific. The tissue-dependent nature of TF-glycogene binding thus requires further validation through either surveying other datasets or performing experimental validation. Many databases curating ChIP-seq and other omic information could aid such TF-glycogene relationship validation. Some examples include: i. ChIP-Seq databases, such as the Gene Transcription Regulatory Database (GTRD) [35]☐ that have analyzed vast publicly available ChIP-Seq data to systematically cataloged TF-gene relationships across several organisms and cellular contexts using multiple ChIP-seq peak calling algorithms, ii. the Regulatory Circuits database [36]☐ that quantifies the activity of promoter and enhancer regions through cap analysis of gene expression (CAGE) and expression Quantitative Trait Loci (eQTL), respectively, and iii. integration of TF binding motifs, protein-protein interactions, and co-expression networks, as was done using data from GTEx and a method called PANDAS [37]. Such cross-platform analysis may reduce the number of candidate TF-glycogene interactions presented in this manuscript. This would also aid more focused wet-lab validation using either CRISRP-Cas9 knockouts and single cell RNA-Seq, or using structural methods that involve glycomics and glycoproteomics based mass spectrometry.

## Conclusion

A majority of current studies in the Glycoscience field use experimental data and curations related to glycans only. Fewer investigations examine the links between the glycans, glycogenes and glycosylation pathways, and other non-glyco datasets. We set out to establish these relationships using bioinformatics and data-mining. Using this, we describe putative regulatory relationships between TFs and glycogenes across 29 cancer types. Some TFs appear to regulate glycogenes in communities, indicating potential cross-talk across pathways in regulating glycosylation. The communities varied with cancer type, even in a single tissue, suggesting that these TF-glycogene interactions are dynamic in nature. Groups of TFs enriched to glycosylation pathways were also associated with signaling pathways, thus establishing a framework to discover how cell signaling processes may regulate cellular glycosylation by modifying TF activity. Overall, the putative TF-glycosylation pathway enrichments found here represent the starting point for wet-lab validation. Such studies could enhance our fundamental understanding of glycosylation pathway regulation, and lead to novel ways to control the glycogenes and glycan structures during disease.

## EXPERIMENTAL METHODS

### Glycogene-pathway classification

A list of 208 unique glycogenes involved in 20 different glycosylation pathways were used in this work (**Supplemental Table S3**). These data are collated from GlycoEnzDB (virtualglycome.org/GlycoEnzDB), with original data coming from various sources in literature [38,39]. The following is a summary of the pathways studied and the enzymes involved:

*<u>1. Glycolipid core:</u>* The enzymes in this group are involved in the biosynthesis of the glucosyl-ceramide (GlcCer) and galactosyl (GalCer)-ceramide lipid core. Here, the GlcCer core is formed by the UDP-glucose:ceramide glucosyltransferase (UGCG) which transfers the first glucose. Following this, lactosylceramide is formed by the action of the β1,4GalT activity of B4GalT5 (and possibly also B4GalT3, 4 and 6). The GalCer core is typically structurally small and is made by UDP-Gal:ceramide galactosyltransferase (UGT8). These structures can be further sulfated by GAL3ST1 or sialylated by ST3GAL5.
*<u>2. P1-Pk Blood Group:</u>* The Pk, P1 and P antigens are synthesized on lactosyl-ceramide glycolipid core. The activity of α1-4GalT (A4GALT) on this core results in the Pk antigen, followed by β1-3GalNAcT (B3GALNT1) to form the P antigen. The P1 antigen, on the other hand, is formed by the sequential action of β1-3GlcNAcT (B3GNT5), β1-4GalT (B4GALT1-6) and α1-4GalT (A4GALT) on the glycolipid core.
*<u>3. Gangliosides:</u>* This pathway encompasses all glycogenes responsible for synthesizing a/b/c gangliosides. UGCG is included to consider the addition of glucose to ceramide. ST3GAL5, and ST8SIA enzymes are added to take the core ganglioside structures to the a,b and c levels. B4GALTs and B4GALNT1 are included to account for ganglioside elongation. Decoration of the gangliosides with sialic acid occurs using ST6GALNAC3-6 and also ST8SIA1/3/5.
*<u>4. Dolichol Pathway:</u>* This results in the formation of the dolichol-linked 14-monosaccharide precursor oligosaccharide. This glycan is co-translationally transferred en bloc onto Asn-X-Ser/Thr sites of the newly synthesized protein as it enters the endoplasmic reticulum. The enzymes involved is such synthesis include the ALG (Asparagine-linked N-glycosylation) enzymes, and additional proteins (part of OSTA and OSTB) involved in the transfer of the glycan to the nascent protein.
*<u>5. Complex N-glycans:</u>* This pathway includes glycogenes responsible for processing the N-linked precursor structure emerging from the dolichol pathway into complex structures. Enzymes involved include mannosidases, glucosidases, some enzymes facilitating protein folding and also enzymes that direct acid hydolases to the lysosome.
*<u>6. N-glycan branching:</u>* These glycogenes are responsible for the addition of GlcNAc to processed N-linked glycan structures. These include all the MGAT enzymes.
*<u>7. GalNAc-type O-glycans:</u>* O-linked glycans are attached to serine or threonine (Ser/Thr) on peptides, where GalNAc is the root carbohydrate. This is mediated by a family of about 20 Golgi-resident polypeptide N-acetylgalactosaminyltransferases (ppGalNAcTs or GALNTs). Core 1 structures result from the attachment of β1-3 linked galactose to the core GalNAc using C1GALT1 and its chaperone C1GALT1C1. Core 2 structures then form upon addition of β1-6 linked GlcNAc by GCNT1. Modifications of core-3 and core-4 glycans can occur during disease and thus this classification includes core-3 forming B3GNT6 and core-4 forming GCNT3. Other O-glycan core-types are rare in nature.
*<u>8. Chondroitin Sulfate & Heparan Sulfate Initiation:</u>* Chondroitin and heparan sulfate glycosaminoglycans all have a common core carbohydrate sequence attaching them to their proteins. These are constructed by the activity of specific Xylotransferases (XYLT1, XYLT2), galactosyltransferses B4GALT7 and B3GALT6 that sequentially add two galactose residues to Xylose, and the Glucuronyltransferase B3GAT3 then adds glucuronic acid to the terminal galactose. Also involved in the formation of this core is FAM20B, a kinase that 2-O-phosphorylates Xylose. At this point, the addition of GalNAc to GlcA by CSGALNACT1 & 2 results in the initiation of chondroitin sulfates chains. The attachment of GlcNAc by EXTL3 to the same GlcA results in heparan sulfates.
*<u>9. Chondroitin/dermatan sulfate extension:</u>* Chondroitin sulfates and dermatan sulfates are extended via the addition of GalNAc-GlcA repeat units. This is catalyzed by CSGALNACT1 which is better suited for the initial GalNAc attachment followed by CSGALNACT2 which is preferred for synthesizing disaccharide repeats. CHSY1, CHSY3, CHPF and CHPF2, all exhibit dual β1,3GlcAT and β1,4GlcAT activity. Additional enzymes mediate sulfation. Epimerization of glucuronic acid to iduronic acid by DSE and DSEL results in the conversion of chondroitin sulfates to dermatan sulfates.
*<u>10. Heparan sulfate extension:</u>* EXT1 and EXT2 both have GlcUA and GlcNAc transferase activities and are together responsible for HS chain polymerization. EXTL1-3 are additional enzymes with GlcNAc transferase activity that facilitate heparin sulfate biosynthesis. Additional enzymes that are critical for heparin sulfate function include the HS2/3/6ST sulfotransferases, the GlcA epimerase GLCE and additional enzymes mediating N-sulfation (NDSTs).
*<u>11. Hyaluronan Synthesis:</u>* This pathway consists of the three hyaluronan synthases HAS1-3.
*<u>12. GPI Anchor Extension:</u>* This pathway includes glycogenes responsible for the synthesis of glycosphosphatidylinositol (GPI) anchored proteins in the ER. This involves synthesis of a glycan-lipid precursor that is en bloc transferred to proteins.
*<u>13. O-Mannose:</u>* This is initiated by the addition of mannose to Ser/Thr using POMT1 or POMT2. β1-2 or β1-4 GlcNAc linkages can then be made using POMGNT1 or POMGNT2 to yield M1 or M3 O-linked mannose structures, respectively. MGAT5B can facilitate β1-4 GlcNAc linkage onto the M1 structure to yield the M2 core. Additional carbohydrates typically found on complex N-linked glycan antennae can then attached. In particular, such extensions may be initiated by members of the B4GALT family or B3GALNT2. Specific variants are noted on α-dystroglycans.
*<u>14. O-linked Fucose:</u>* This pathway includes POFUT1, the enzyme responsible for the addition of fucose to Ser/Thr residues. MFNG, LFNG and RFNG which can attach β3GlcNAc to this fucose.
*<u>15. Type 1 & 2 LacNAc:</u>* These enzymes help construct either Galβ1,3GlcNAc (Type 1) or Galβ1,4GlcNAc (Type 2) lactosamine chains on antennae of N-linked glycan, O-linked glycans and glycolipids. Also included are GCNT1-3 that can facilitate formation of I-branches on N-glycans.
*<u>16. Sialylation:</u>* This group encompasses all kinds of sialyltransferases: ST6GAL, ST3GAL, ST8SIA, and ST6GALNACs. Enrichments to this pathway capture overall increase in sialylation regardless of context.
*<u>17. Fucosylation:</u>* These include α1-2 (FUT1,2) and α1-3 (FUT3, 4, 5, 6, 7, 9) fucosyltransferases that can act on N-glycans, O-glycans and glycolipids.
*<u>18. ABO blood Group Synthesis:</u>* These are enzymes involved in the biosynthesis of ABO antigens
*<u>19. LacDiNAc:</u>* Glycogenes involved in the synthesis of LacDiNac structures.
*<u>20. Sulfated glycan epitopes:</u>* This includes the enzymes attaching sulfate to different types of carbohydrates

### Establishing TF–glycogene relationships in Cistrome Cancer

Regulatory potential and gene correlation data were downloaded from the Cistrome Cancer database in tab-delimited form (http://cistrome.org/CistromeCancer/CancerTarget/)[14]. TF-gene relationships were filtered for the 341 glycogenes in this manuscript (**Supplemental Table S1**). In total, the full dataset contained 41,771 TF-to-glycogene relationships, including relational data for 568 unique TFs found in the 29 cancer systems across all the glycogenes. Positive regulatory relationships between TFs and glycogenes were selected based on RP ≥ 0.5 and ρ ≥ 0.4 (**Fig 2**). This filtering resulted in 20,617 TF-glycogene relationships including 524 unique TFs across 29 cancer types.

Cytoscape was used to visualize TF-glycogene regulatory relationships [40]. To achieve this, all TF-glycogene relationship data were loaded into cytoscape as a network. These data were filtered based on RP and *ρ* thresholds defined previously. A binding potential (BP) score was computed by taking the product of RP and *ρ* for each TF-glycogene relationship. TF-glycogene relationships for each cancer type were separated into sub-networks. The prefuse force directed layout algorithm in cytoscape was used to arrange nodes in each cancer sub-network. The closeness of nodes to one another is weighted by 1-BP. Thus, nodes with high BPs will be placed closer together, whereas smaller BPs will be placed further away. Since there are two classes of nodes (TFs and glycogenes), we treated TF-glycogene networks as bipartite, and applied the corresponding procedure for community detection [20]. Firstly, the bipartite TF-glycogene graphs are projected into two different unipartite graphs, where TFs and glycogenes are placed into separate graphs. The edge weights connecting TFs is computed as the number of shared glycogenes they regulate. The TF unipartite graph was then subjected to a greedy modularity optimization-based approach implemented in the R package igraph [41]. TF-glycogene interactions in each community were subjected to overrepresentation analyses to identify enriched signaling and glycosylation pathways.

### Relating TF-glycogene interactions to glycosylation and signaling pathways

A Fisher’s Exact Test was applied to determine if a particular TF disproportionately regulates one of the 20 glycosylation pathways described in **Supplemental Table S3**. Input data to the test consisted of all TF-gene interactions that passed the RP and ρ thresholds for the cancer type being analyzed. TFs were considered to be disproportionately regulating a glycosylation pathway if the Fisher’s exact test resulted in a p-value≤ 0.05.

TFs enriched to glycosylation pathways were associated with putative regulatory pathways using Reactome DB over-representation analysis API, which uses a Fisher’s exact test, to associate the TFs with signaling pathways[12]. Signaling pathway enrichments with adjusted p (FDR)<0.1 were kept. A high p-value cutoff was chosen to allow users to gain a high-level perspective as to what potential pathways may be regulating enriched TFs. The connection between cell signaling pathways and TFs, and that between the TFs and glycosylation pathways were visualized using alluvial plots generated using the R package ggalluvial. Only signaling pathways with < 30 members are presented for brevity. A comprehensive listing of enriched signaling pathways is available in Supplemental Table S5.

## SUPPORTING INFORMATION

Supplementary Table S1:

File Name: TableS1_Glycogenes.xlsx

File Format: XLXS

Title: List of 341 glycogenes used for cystoscape maps

Supplementary Table S2:

File Name: TableS2_CancerTypes.xlsx

File Format: XLSX

Title: Cancer type list

Supplementary Table S3:

File Name: TableS1_Glycogene_Pathway_Lists.xlsx

File Format: XLSX

Title: Glycogene pathway lists

Supplementary Table S4:

File Name: TableS4_Fishers_exact_test_summary.xlsx

File Format: XLXS

Title: Fisher’s exact test to infer TF-glycosylation pathway relation (p<0.05 data are highlighted)

Supplementary Table S5:

File Name: TableS5_Reactome_Enrich_Pathways.xlsx

File Format: XLXS

Title: Reactome pathway enrichments for all TFs

Supplementary Table S6:

File Name: TableS6_GlycoPathway_Network_Enrichment.xlsc

File Format: XLXS

Title: Network Cluster Enriched Glycosylation Pathways

File Name: TableS7_Signaling_Network_Enrichment. xls

File Format: XLXS

Title: Network Cluster Enriched Signaling Pathways from Reactome

Supplementary File S1:

File Name : FileS1_supplemental_alluvials.docx

File Format: docx

Title: Alluvial plots for all cancer types

Supplementary File S2:

File Name: FileS2_CancerNetworks_Cistrome.cys

File Format : cys (Cytoscape Session File)

Title: Cistrome Cancer TF-to-glycogene subnetworks

R scripts used for analysis are available at: https://github.com/neel-lab/TF-glycogene

## ACKNOWLEDGEMENT

We gratefully acknowledge helpful discussions with Prof. Rudiyanto Gunawan

