## Supplementary figures and images for "A systems based framework to computationally predict putative transcription factors and signaling pathways regulating glycan biosynthesis"

### FigureS1_Alluvials_V2.docx

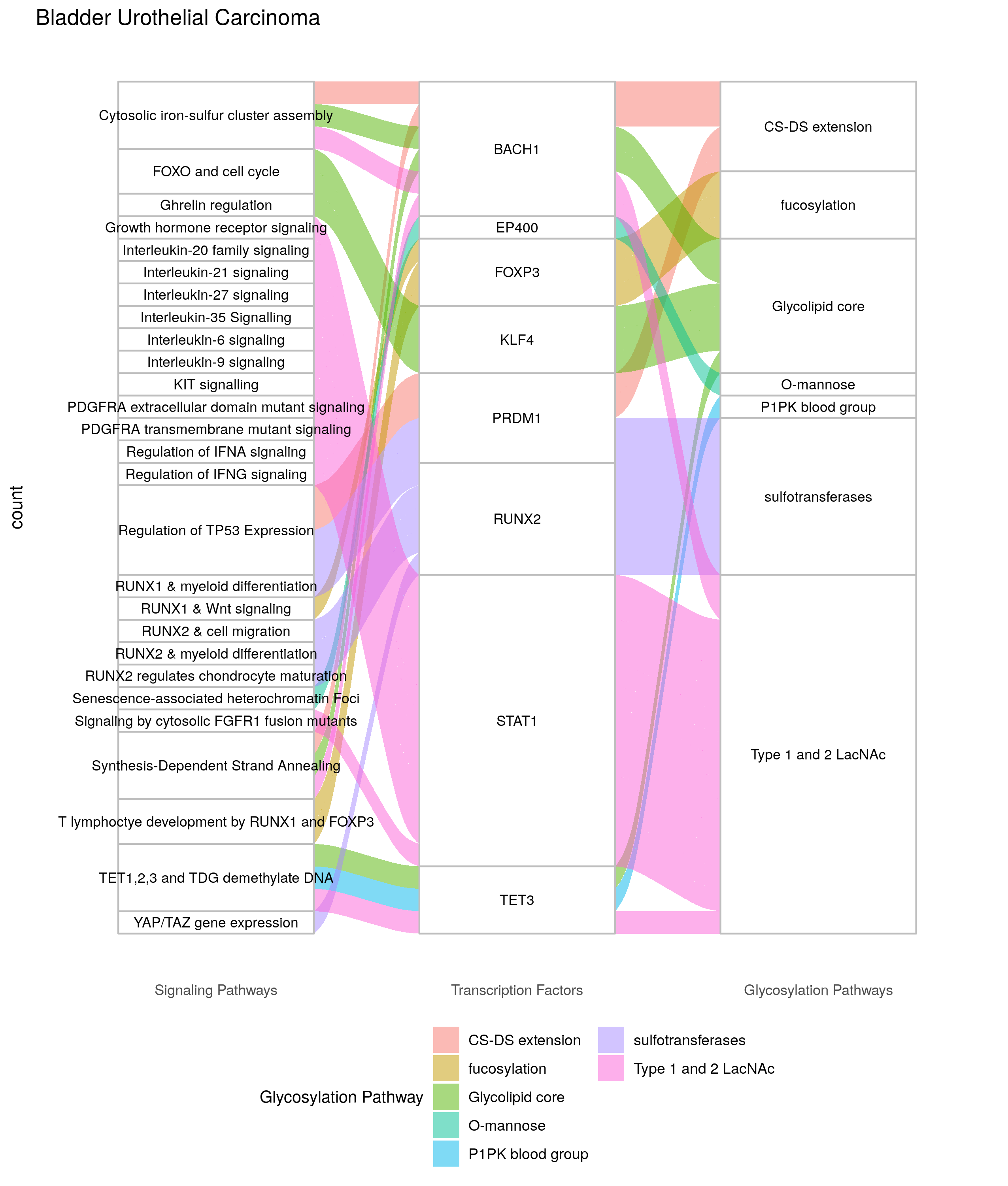

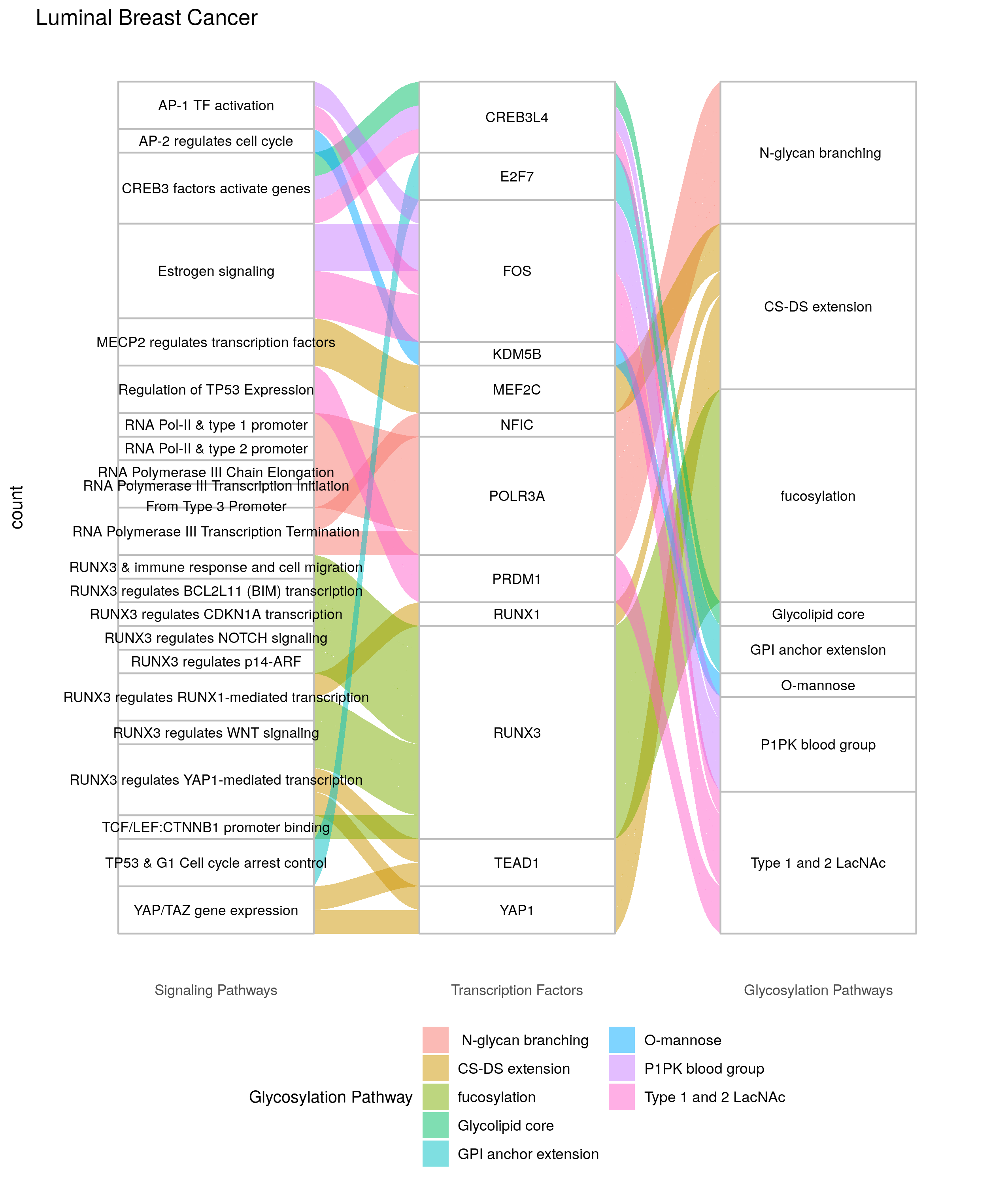

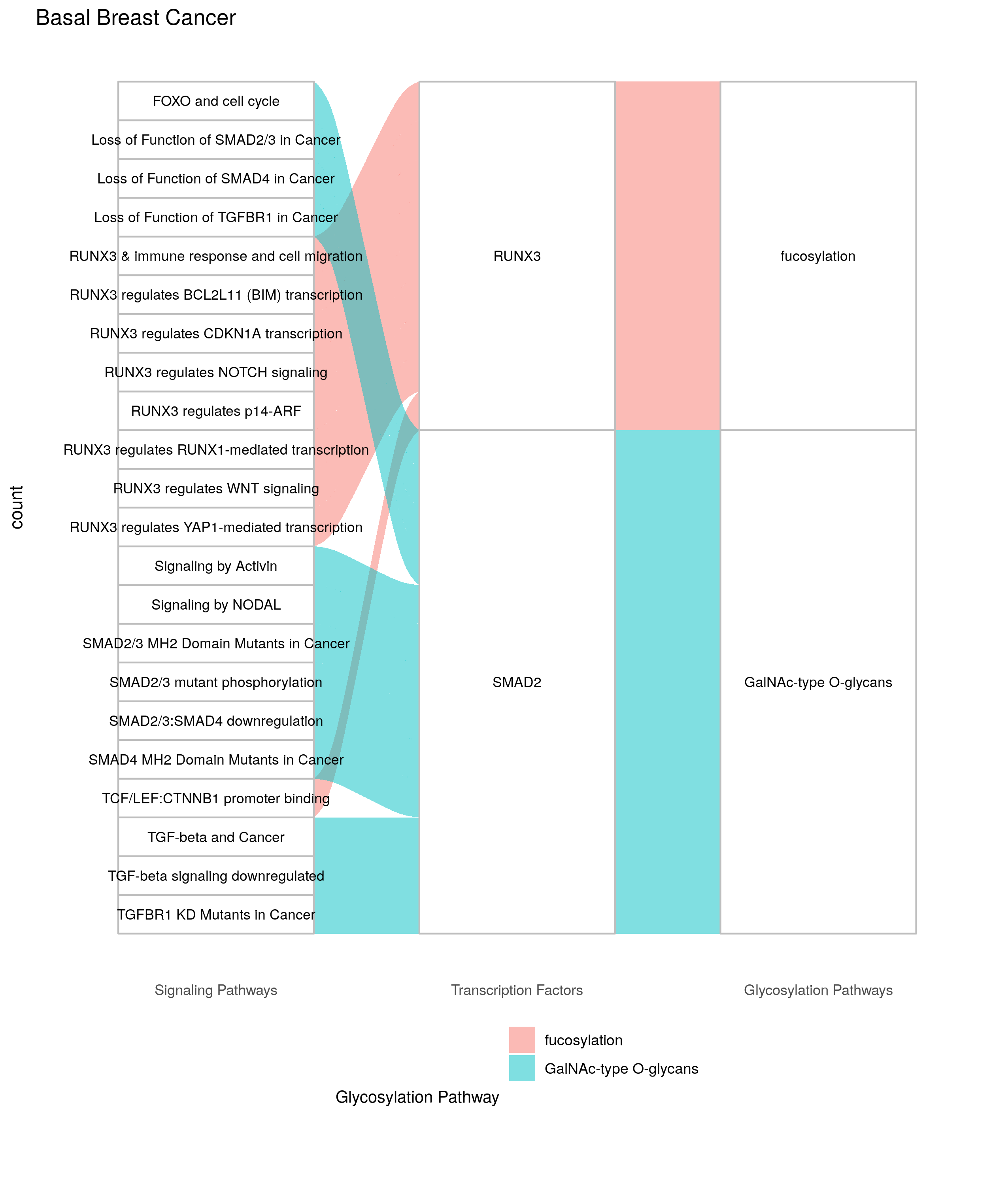

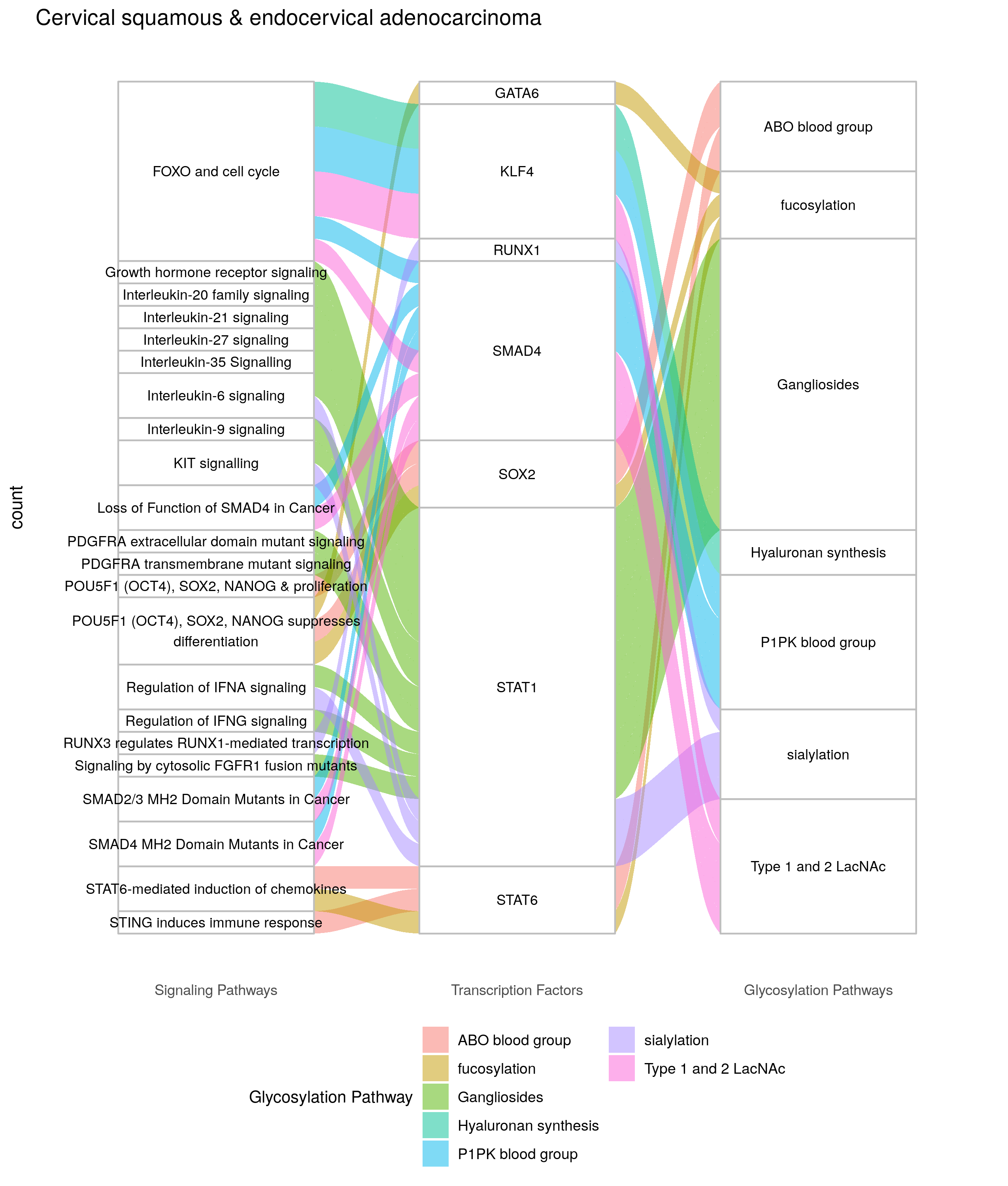

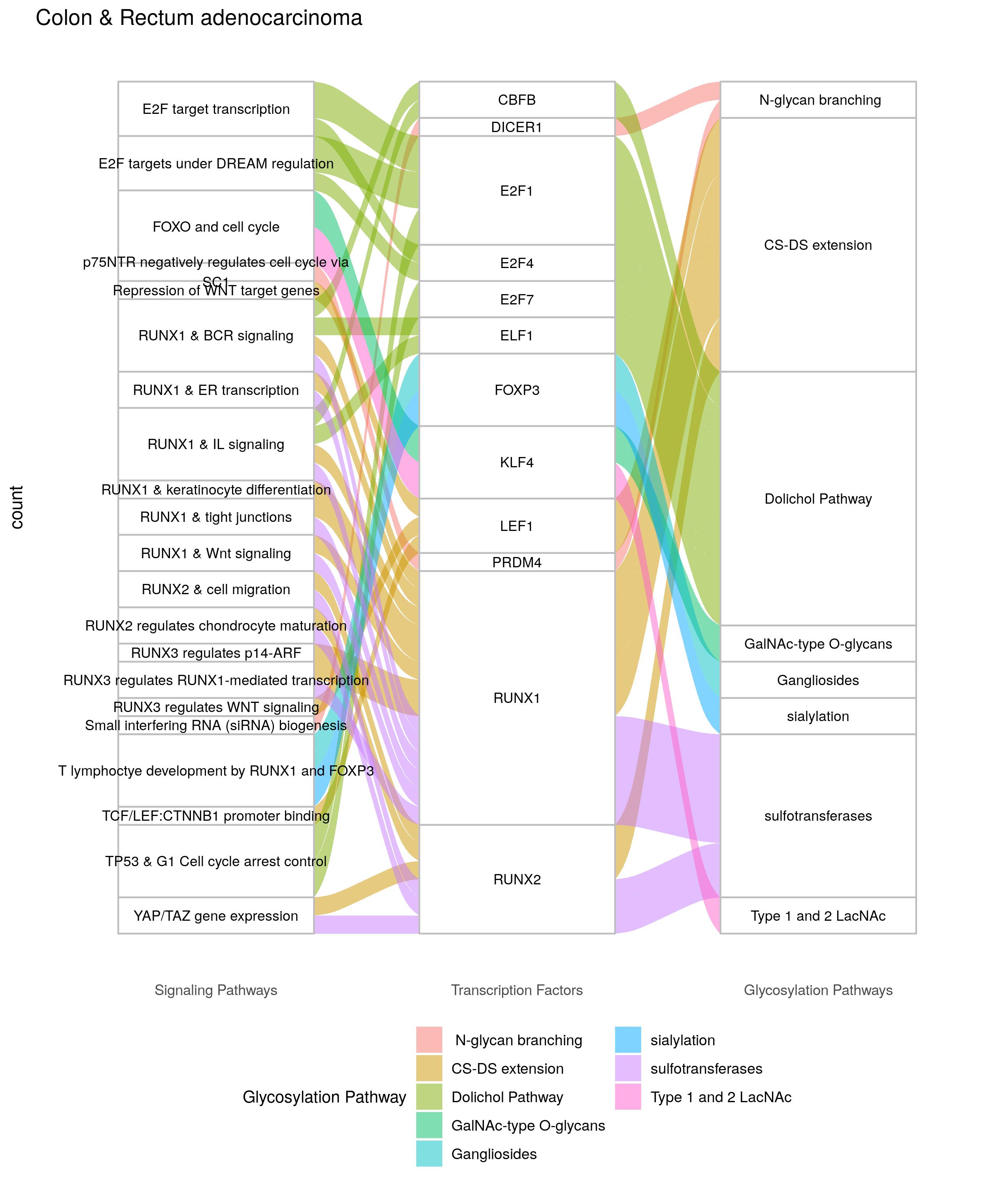

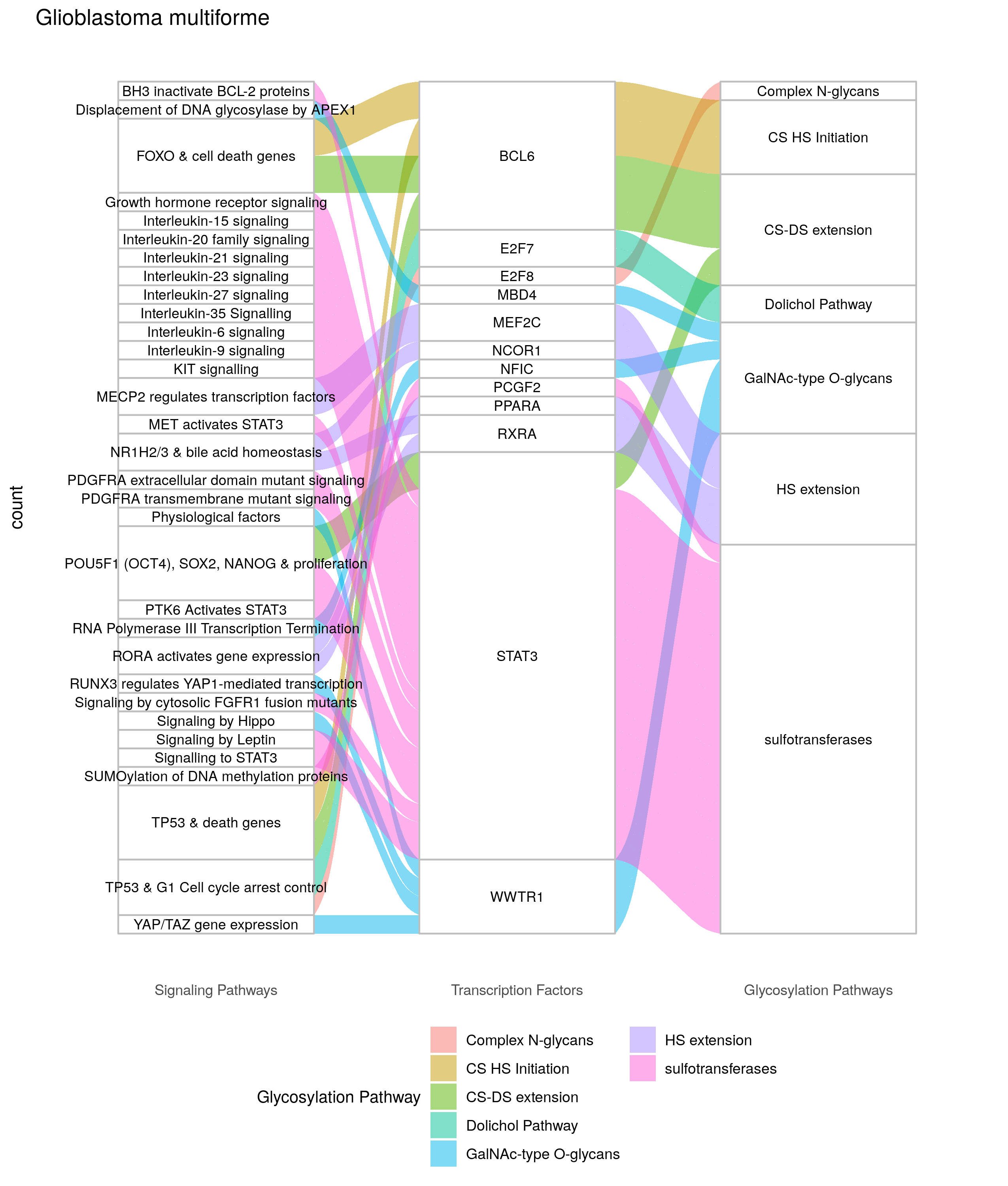

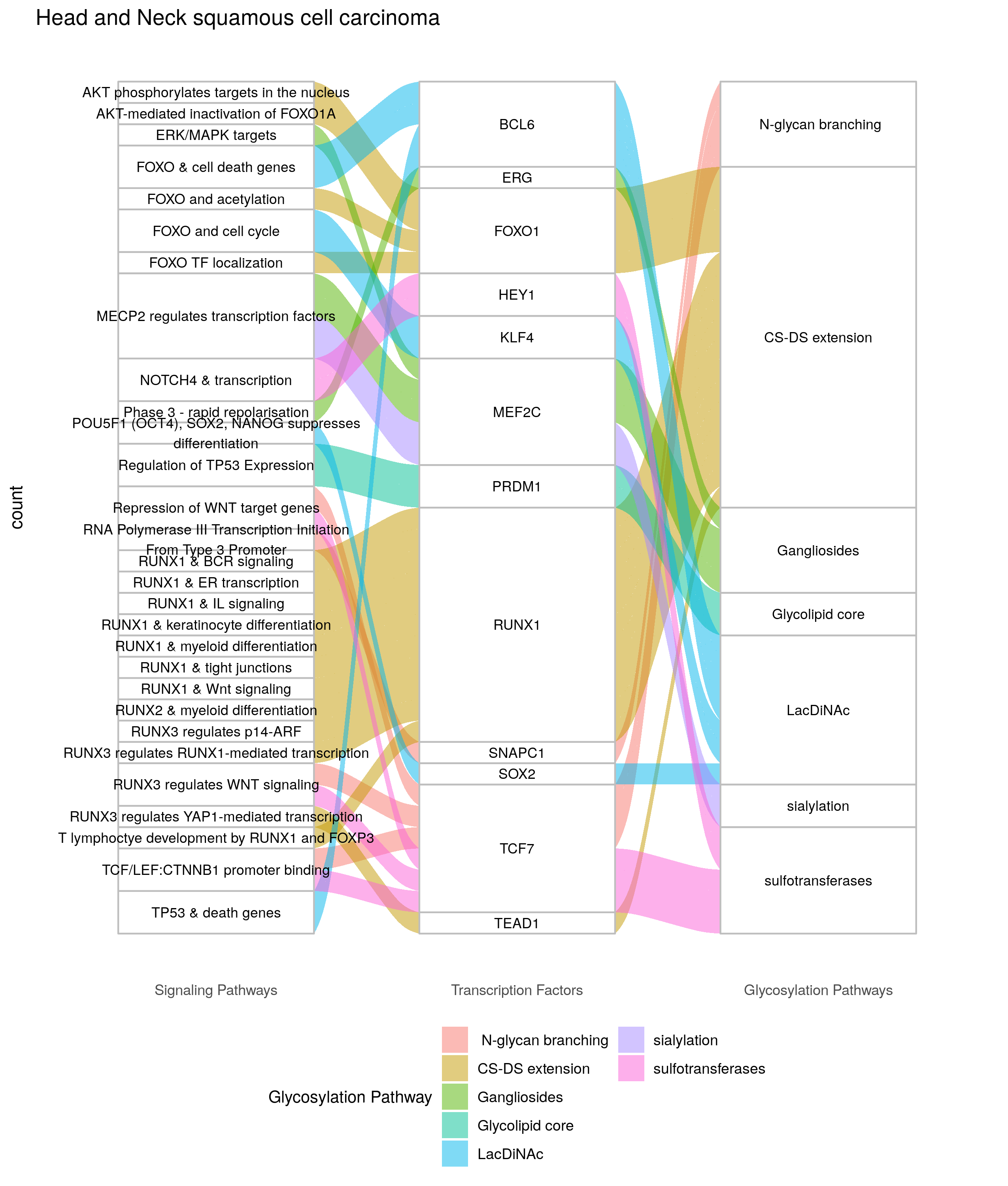

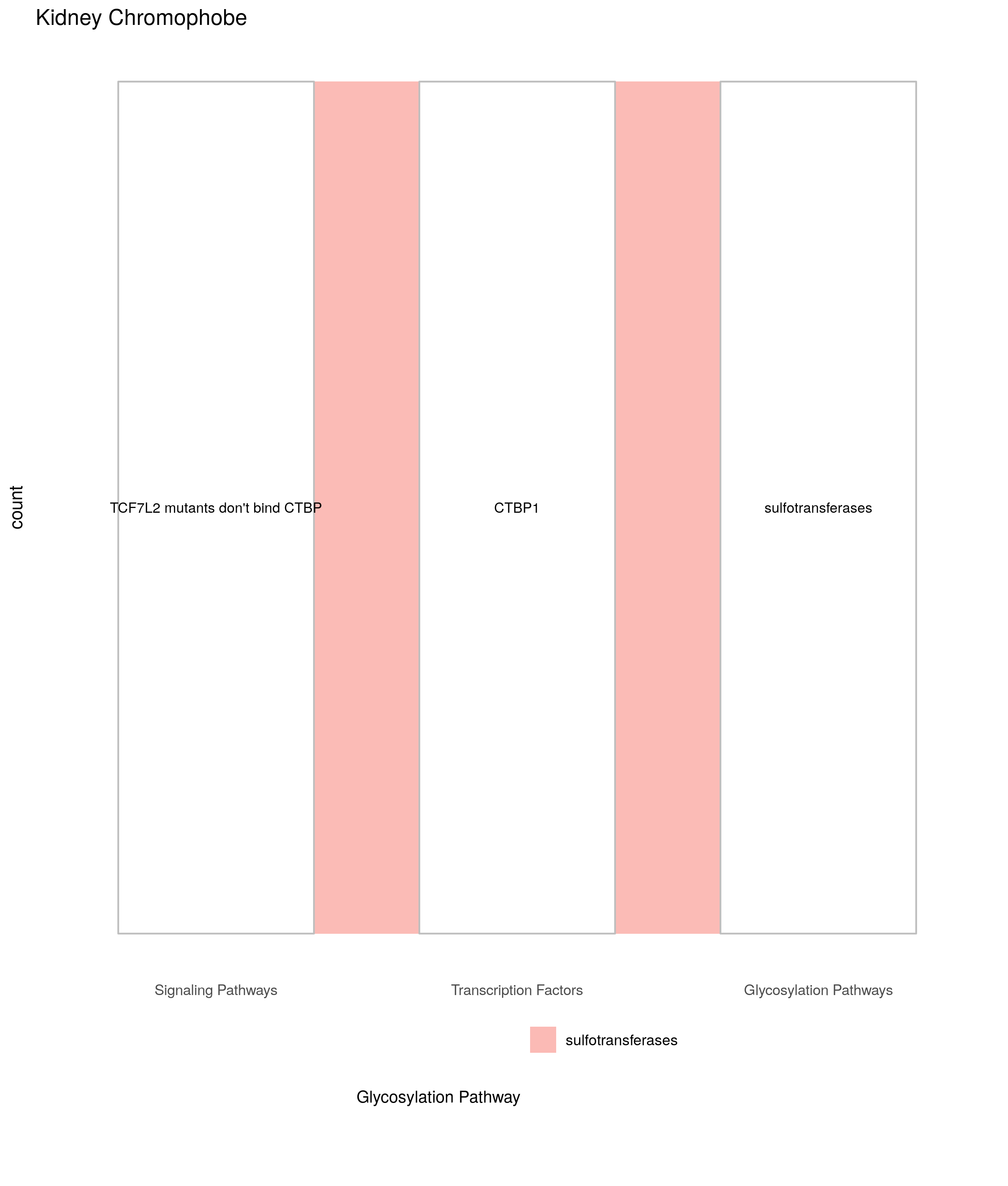

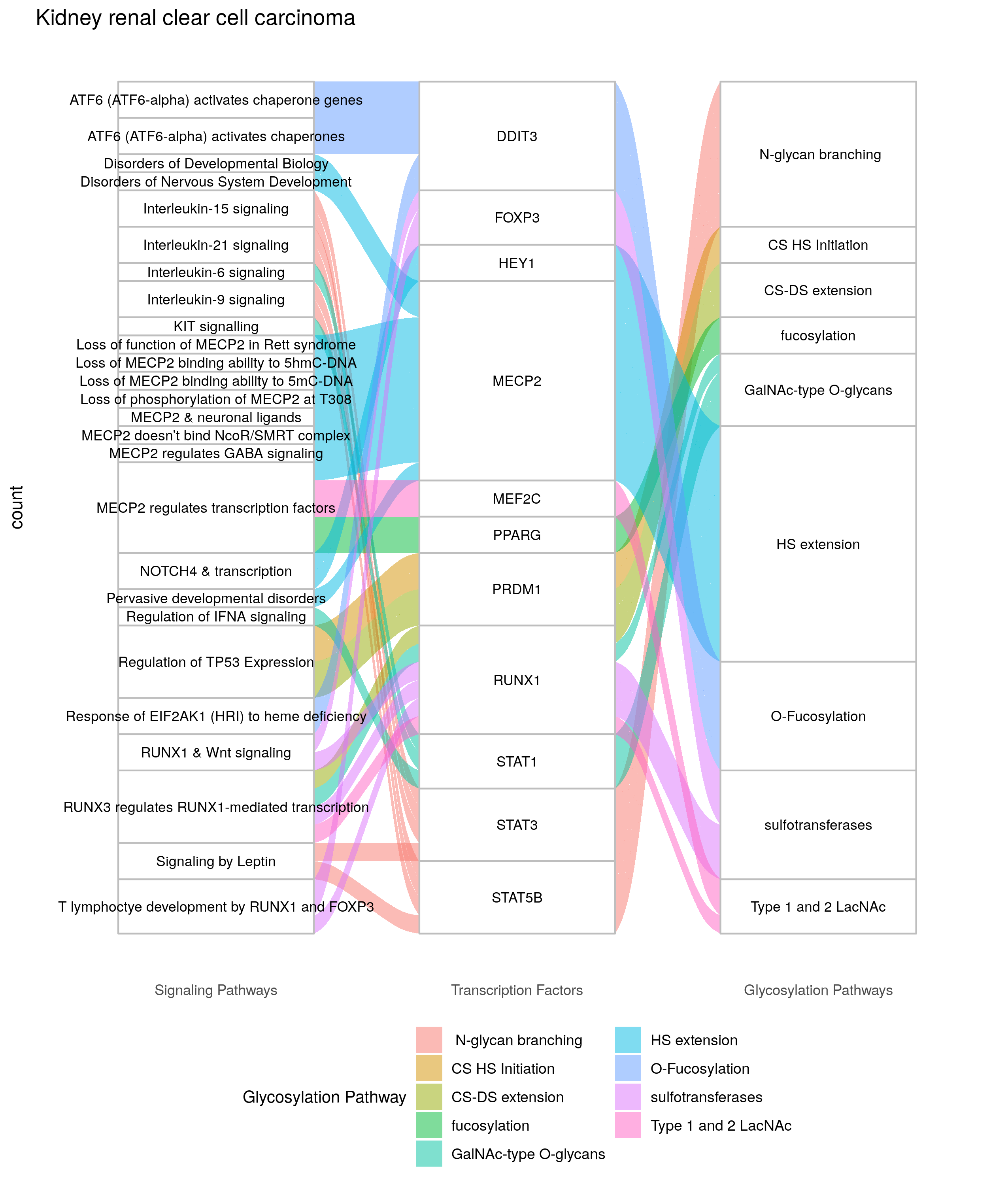

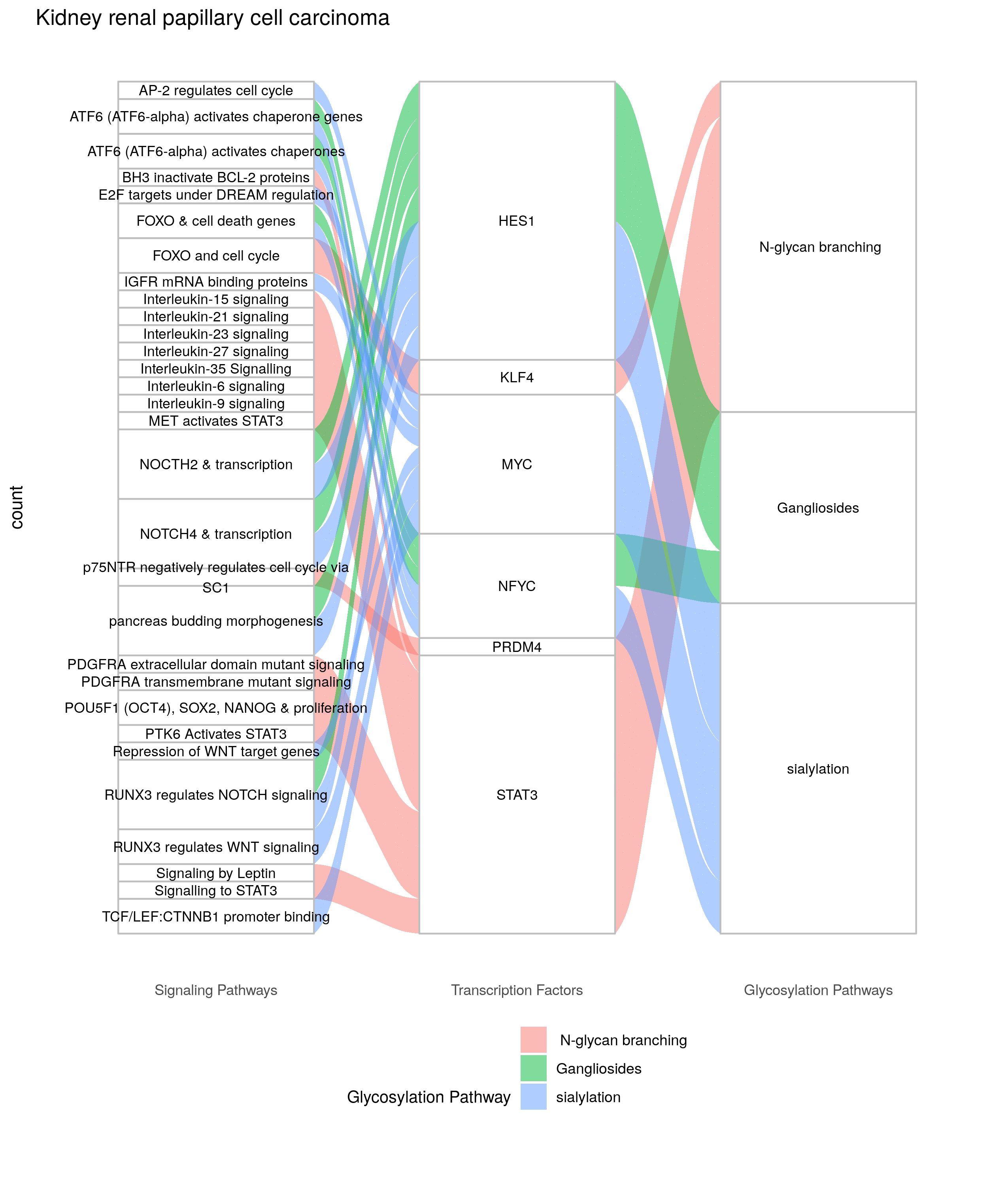

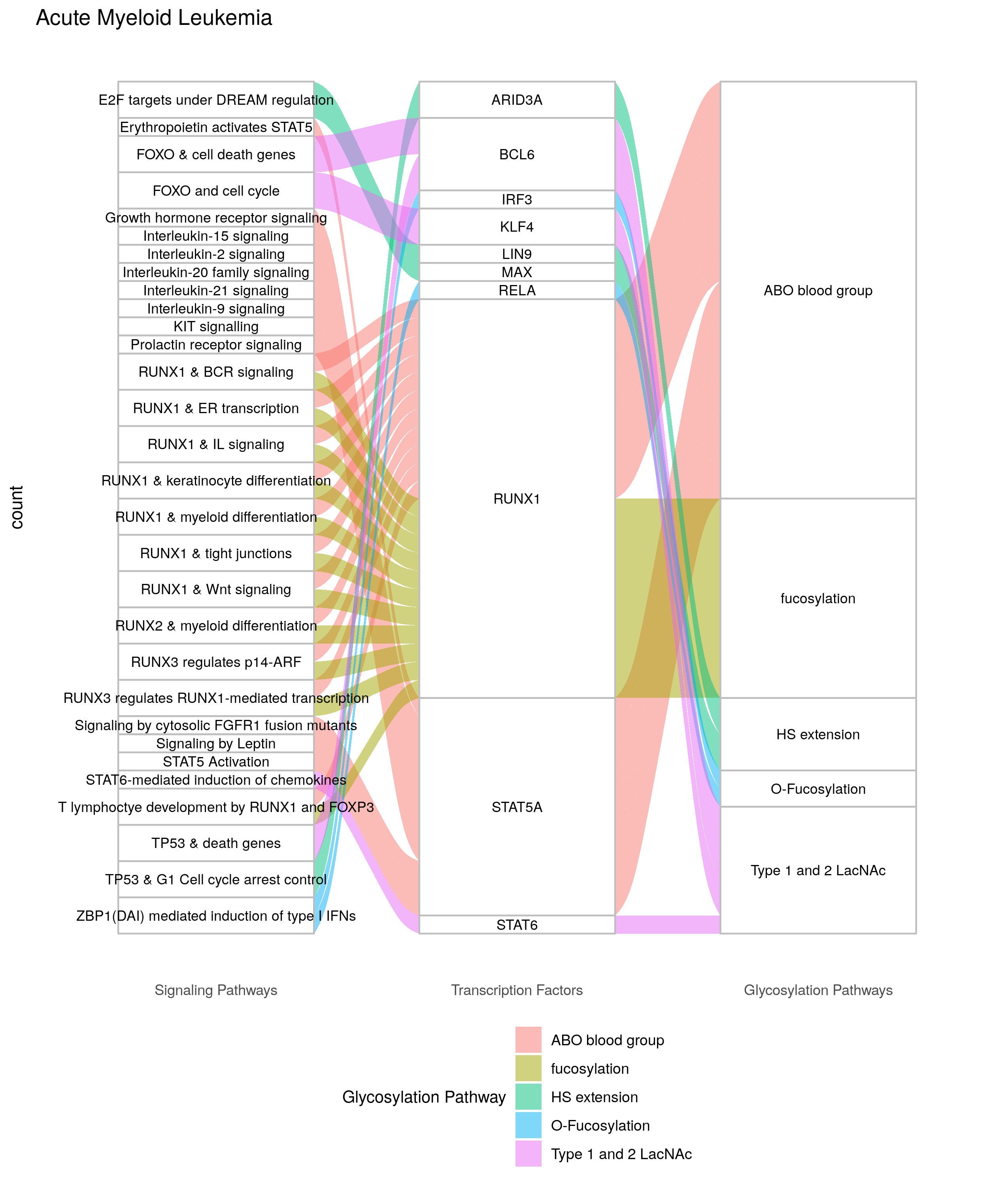

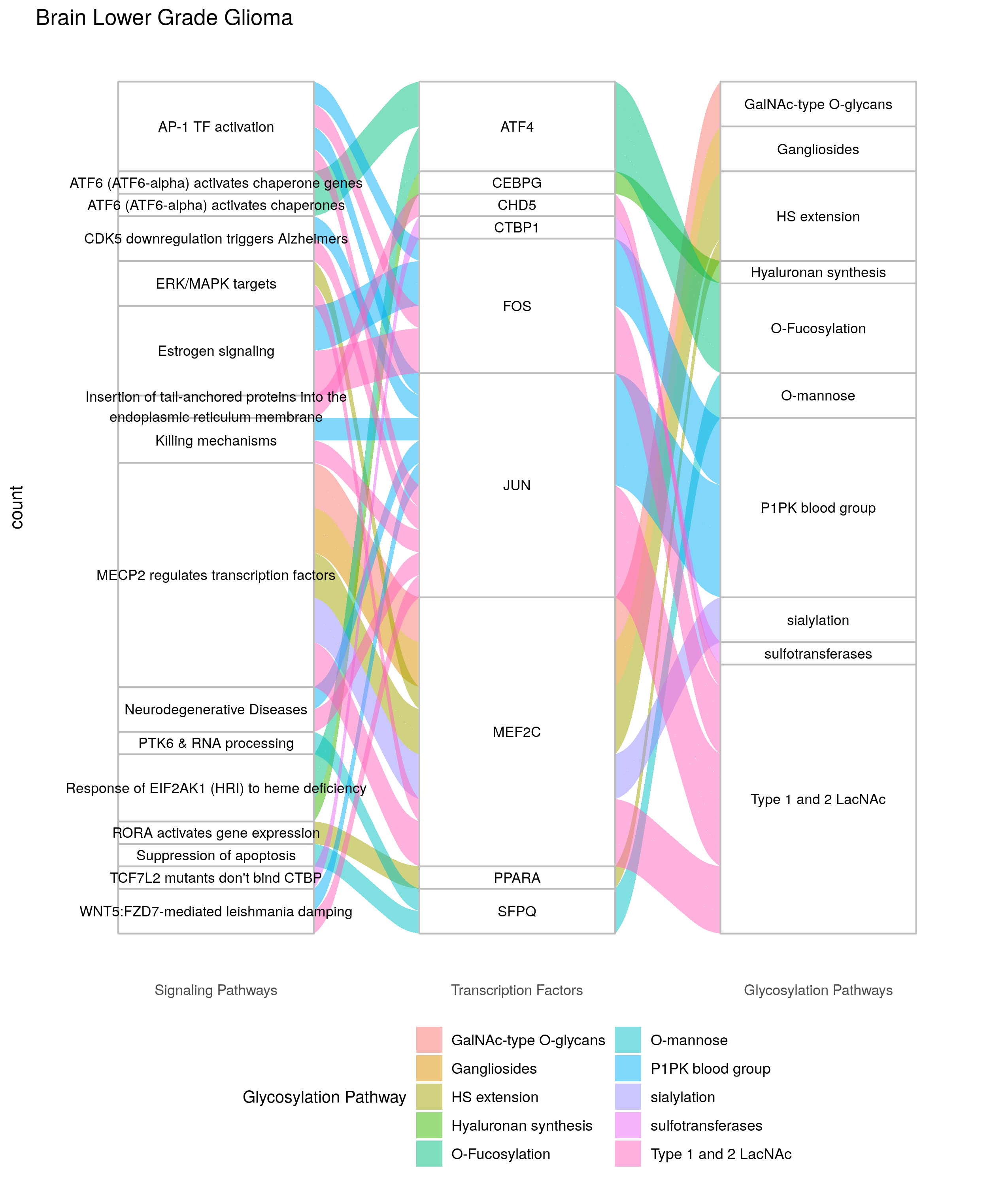

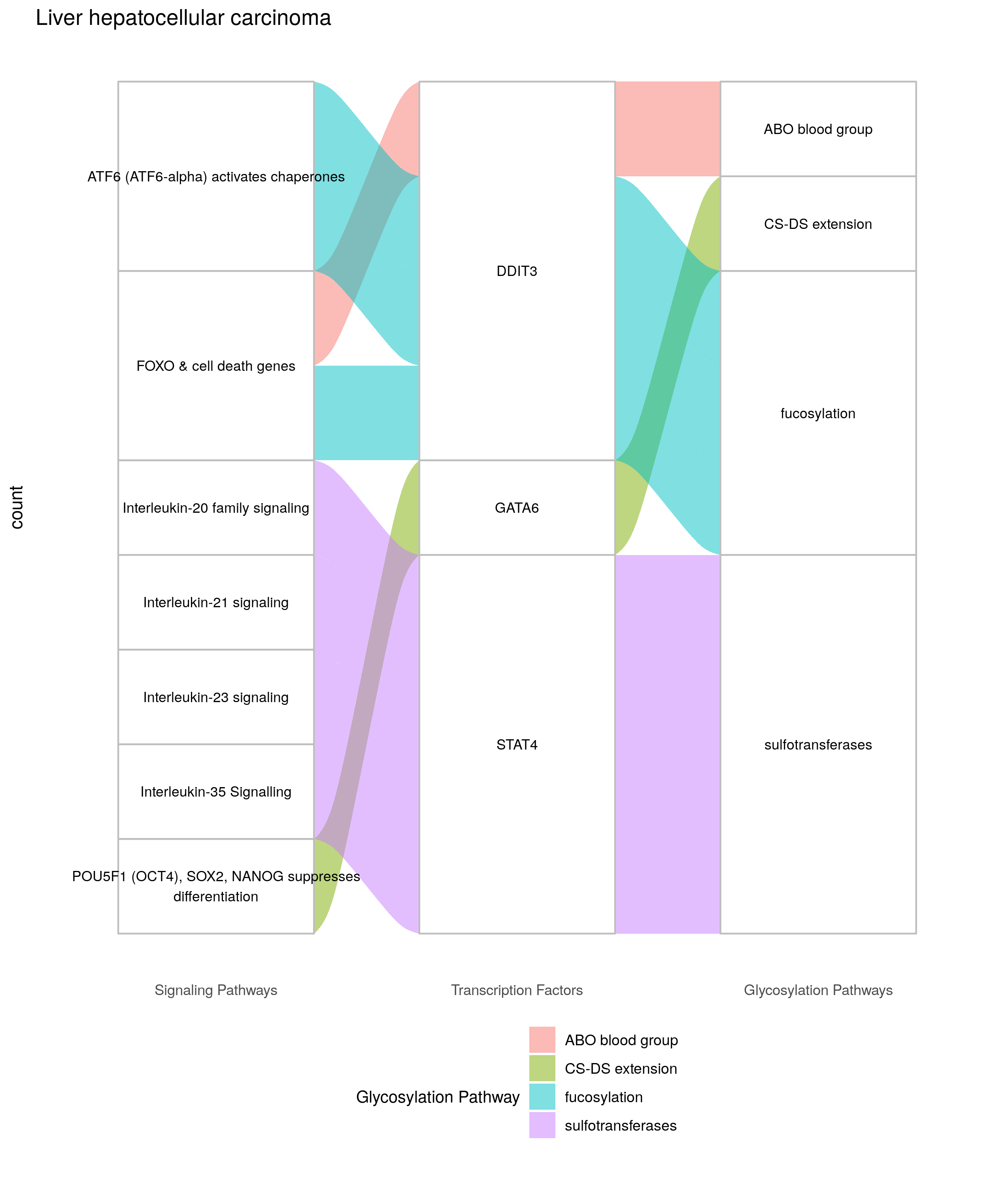

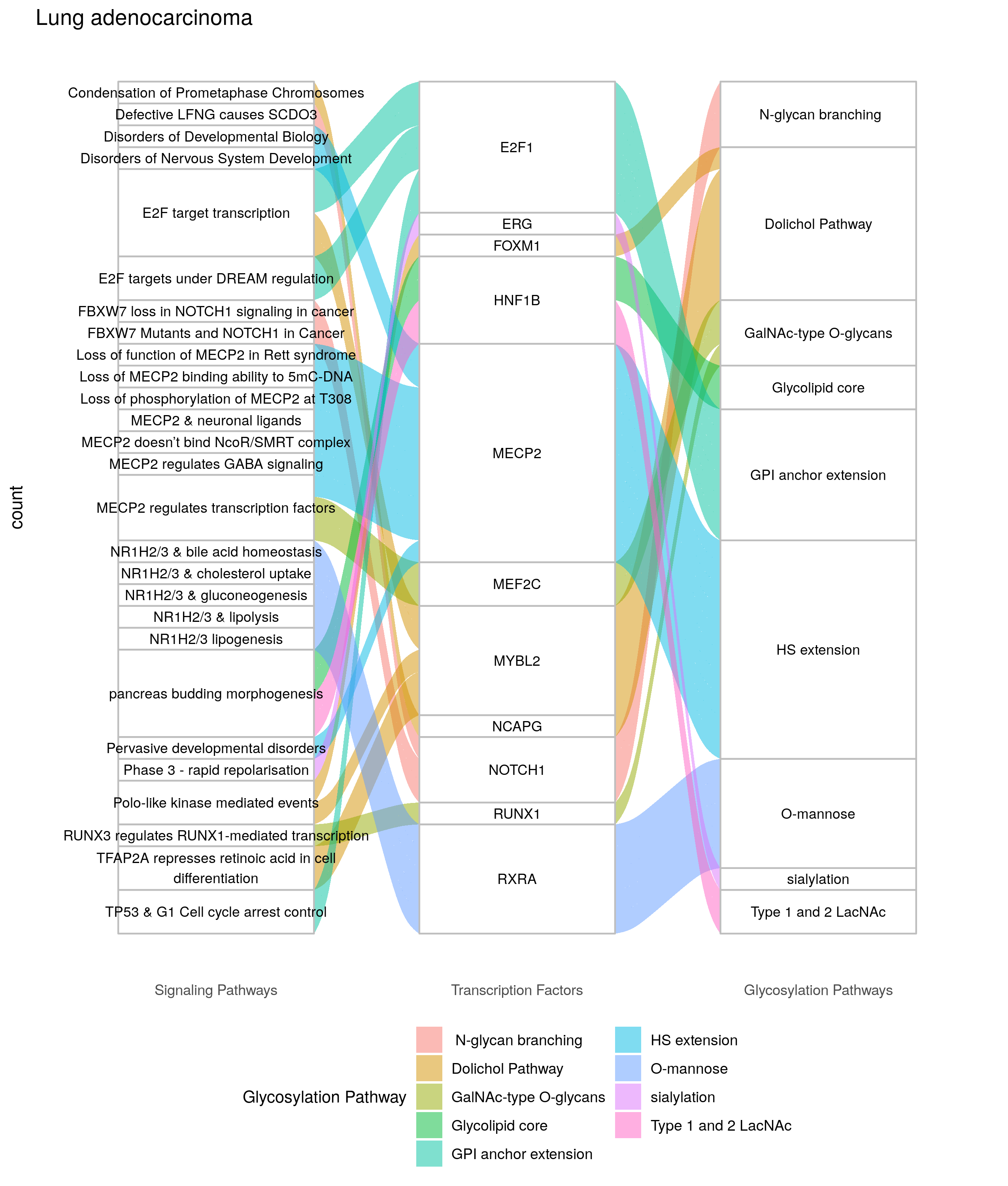

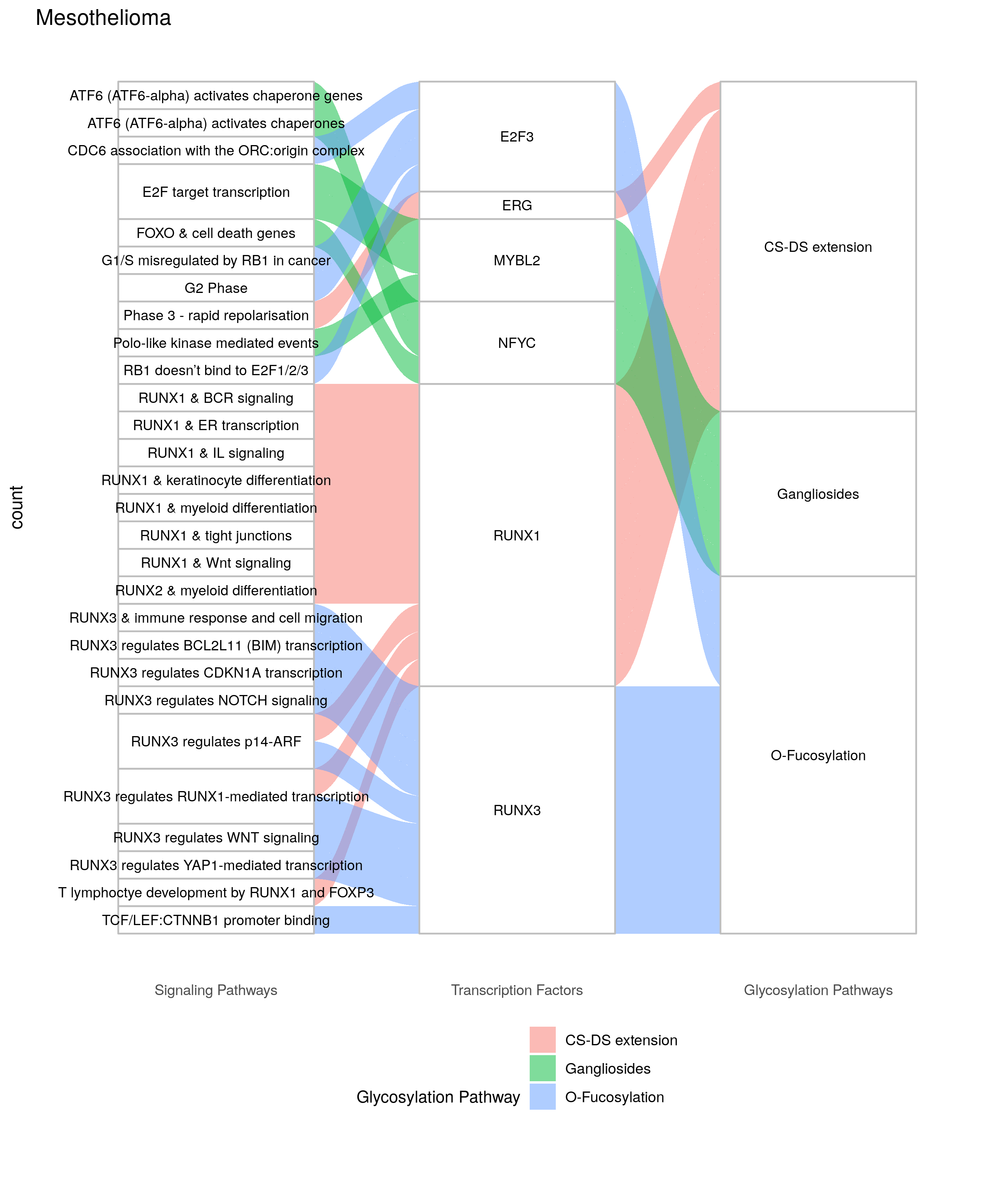

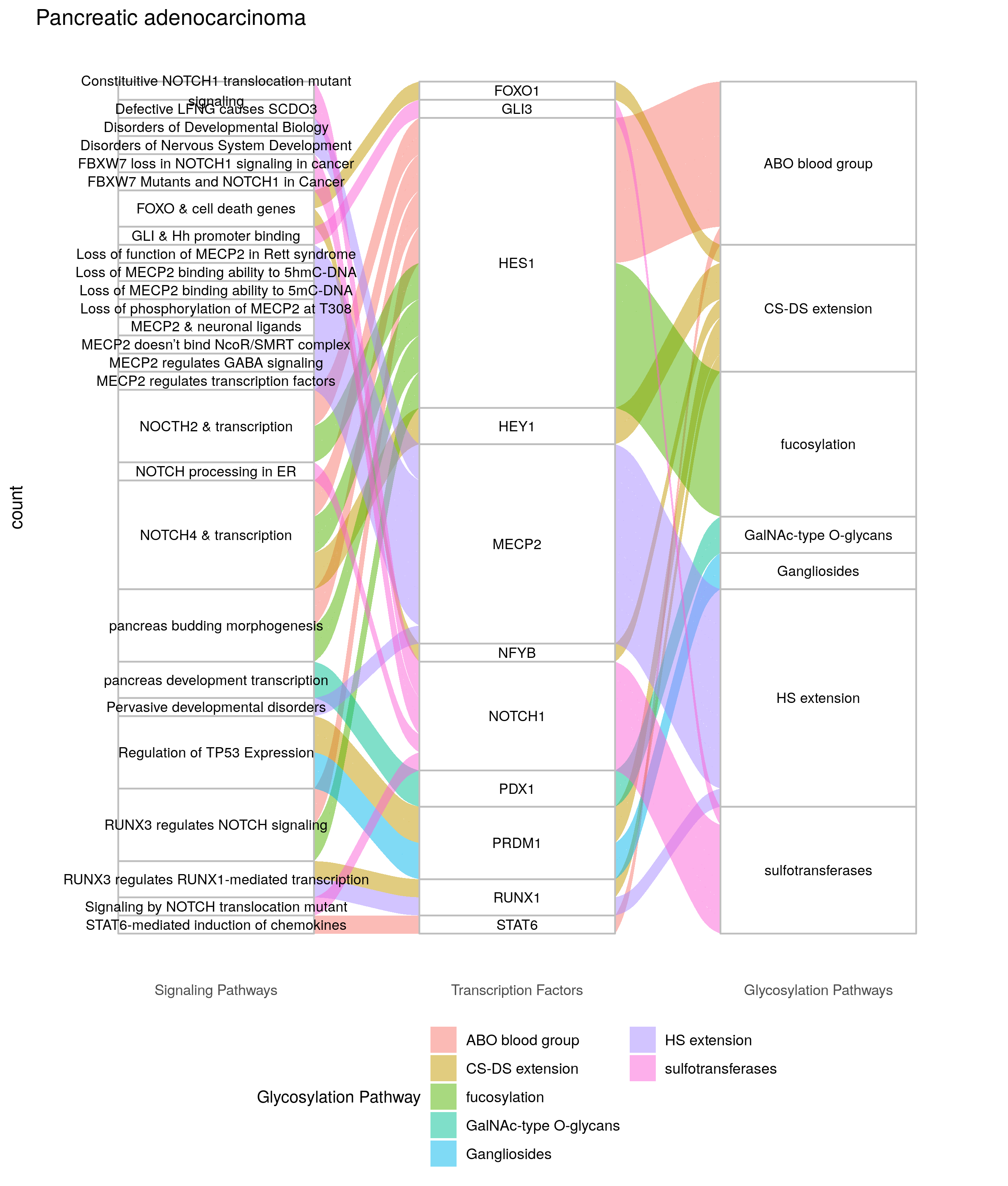

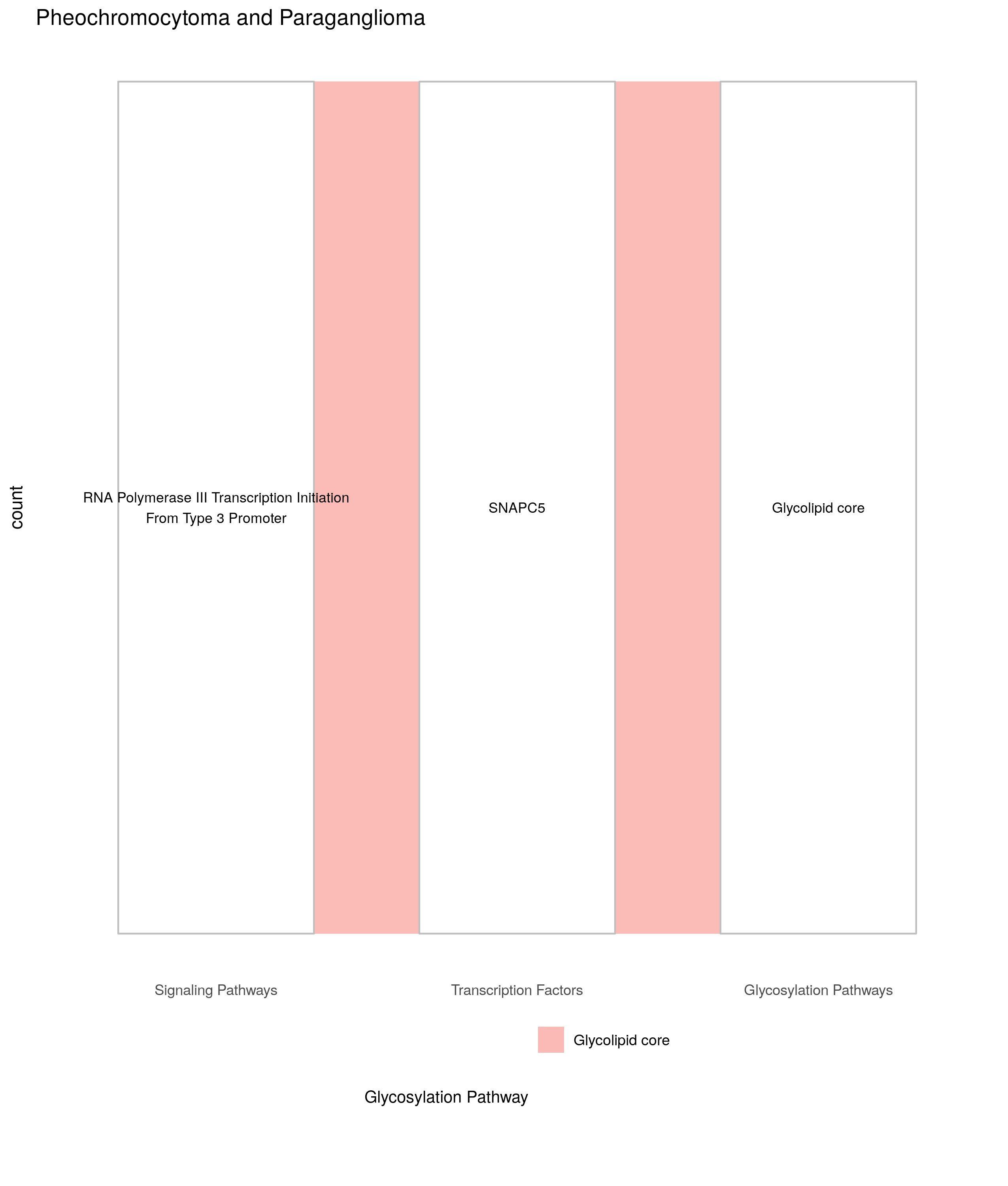

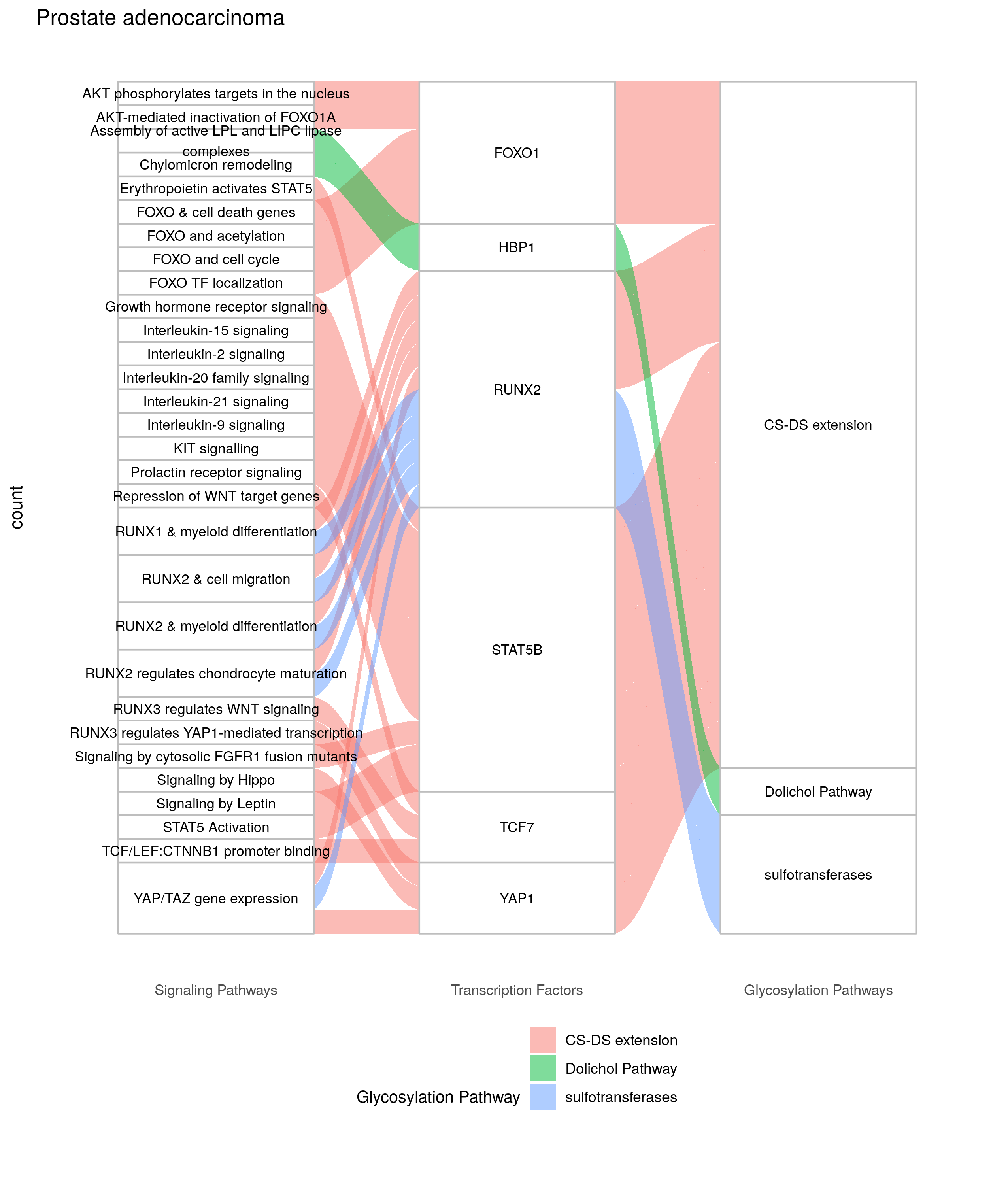

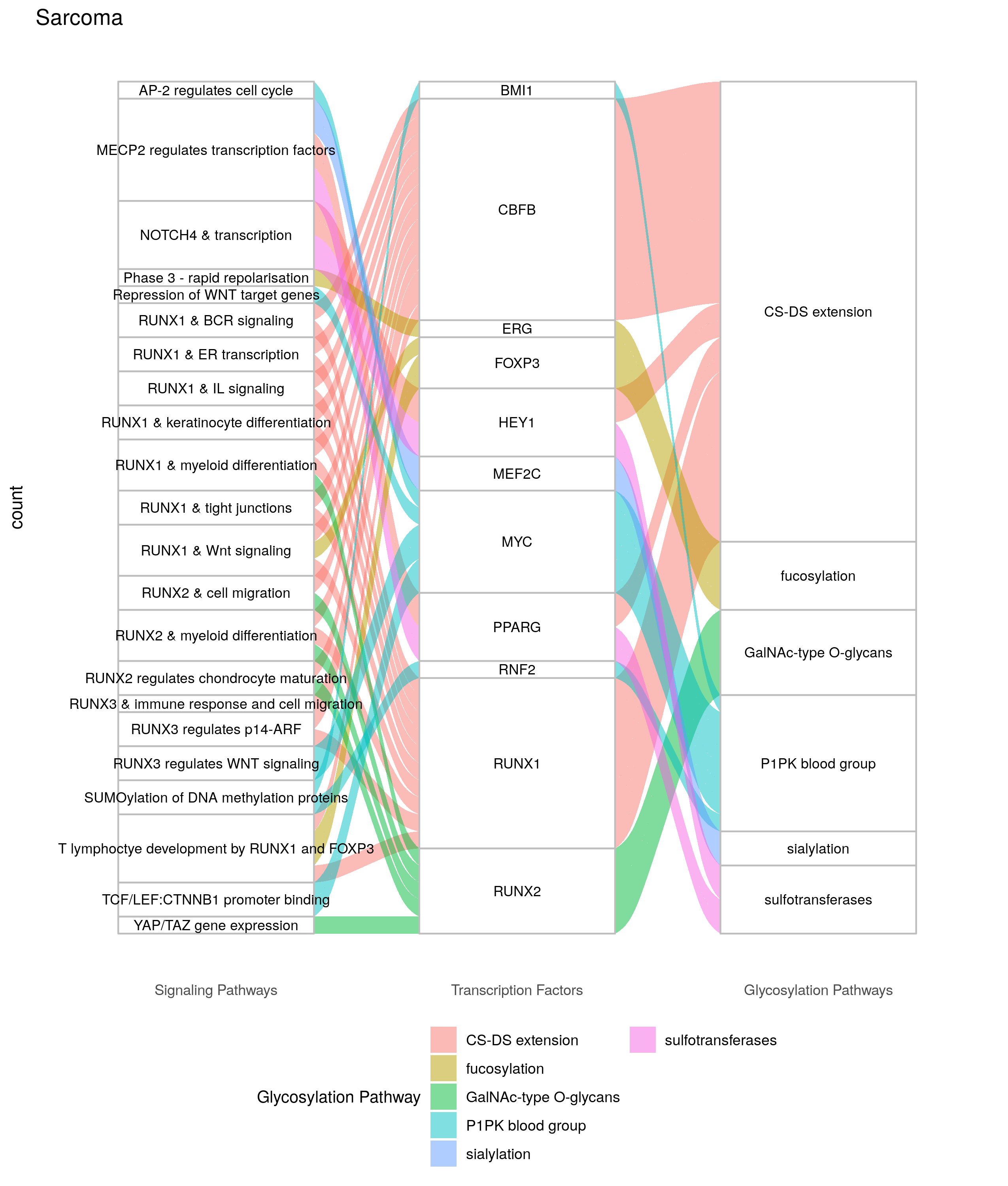

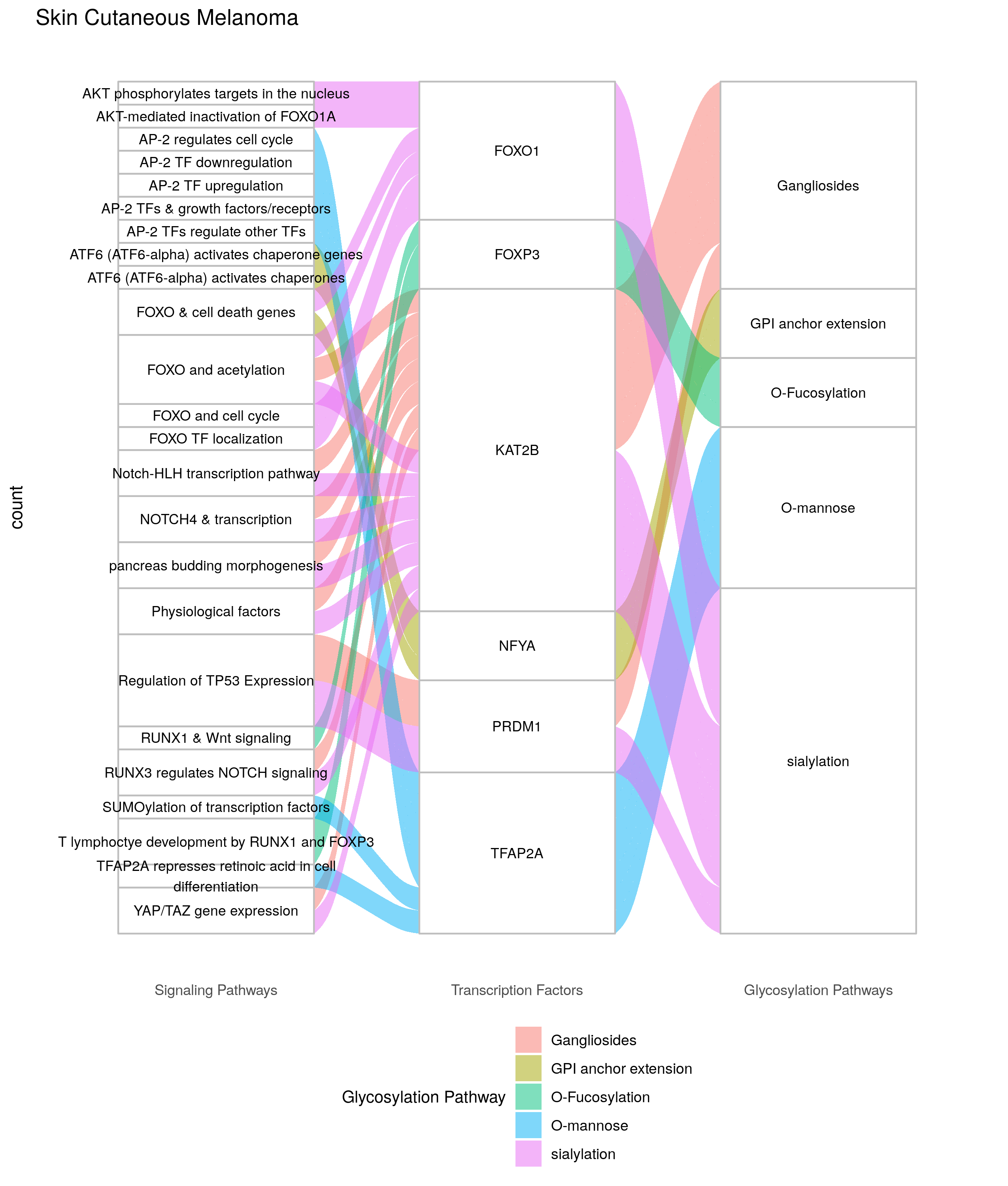

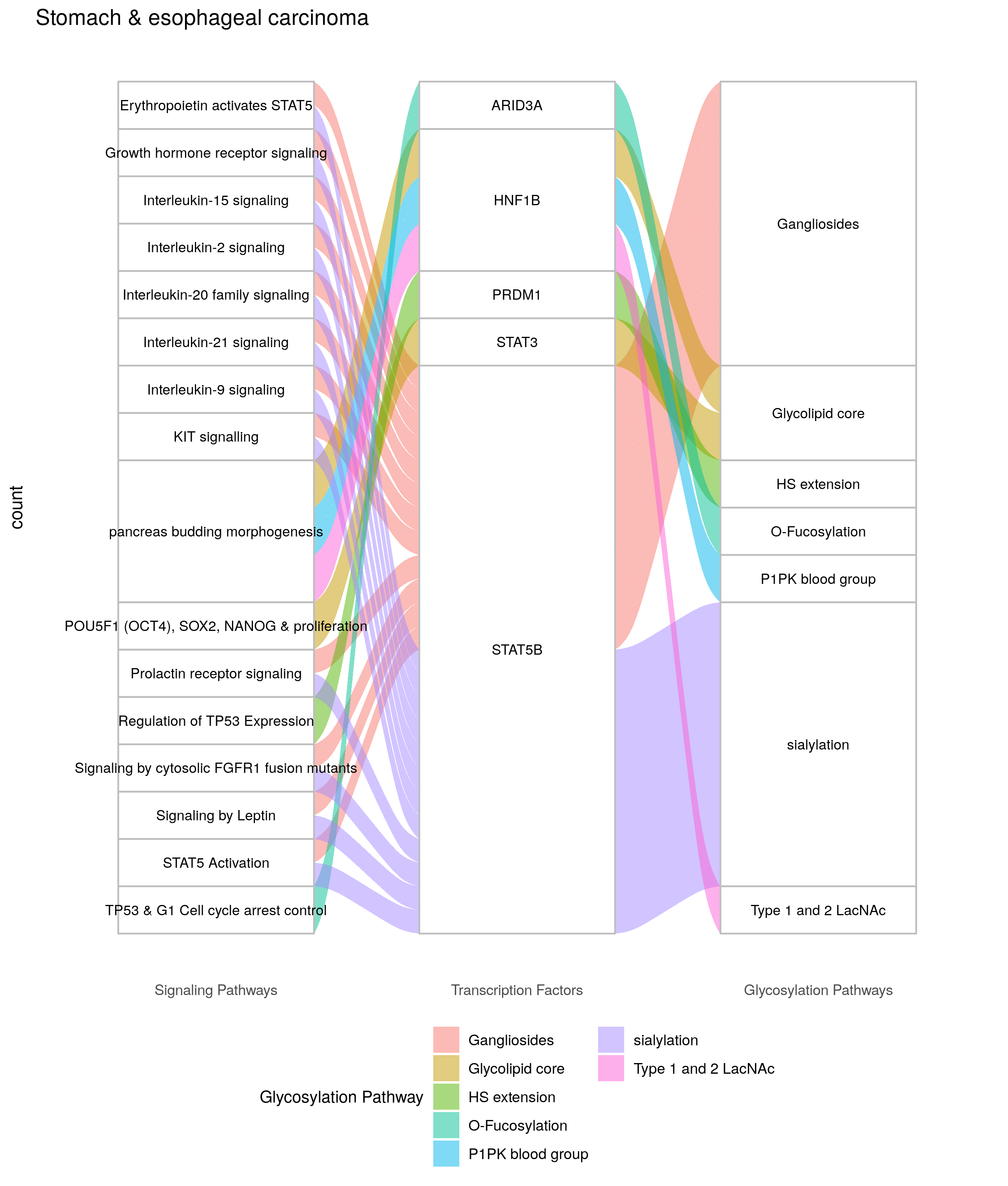

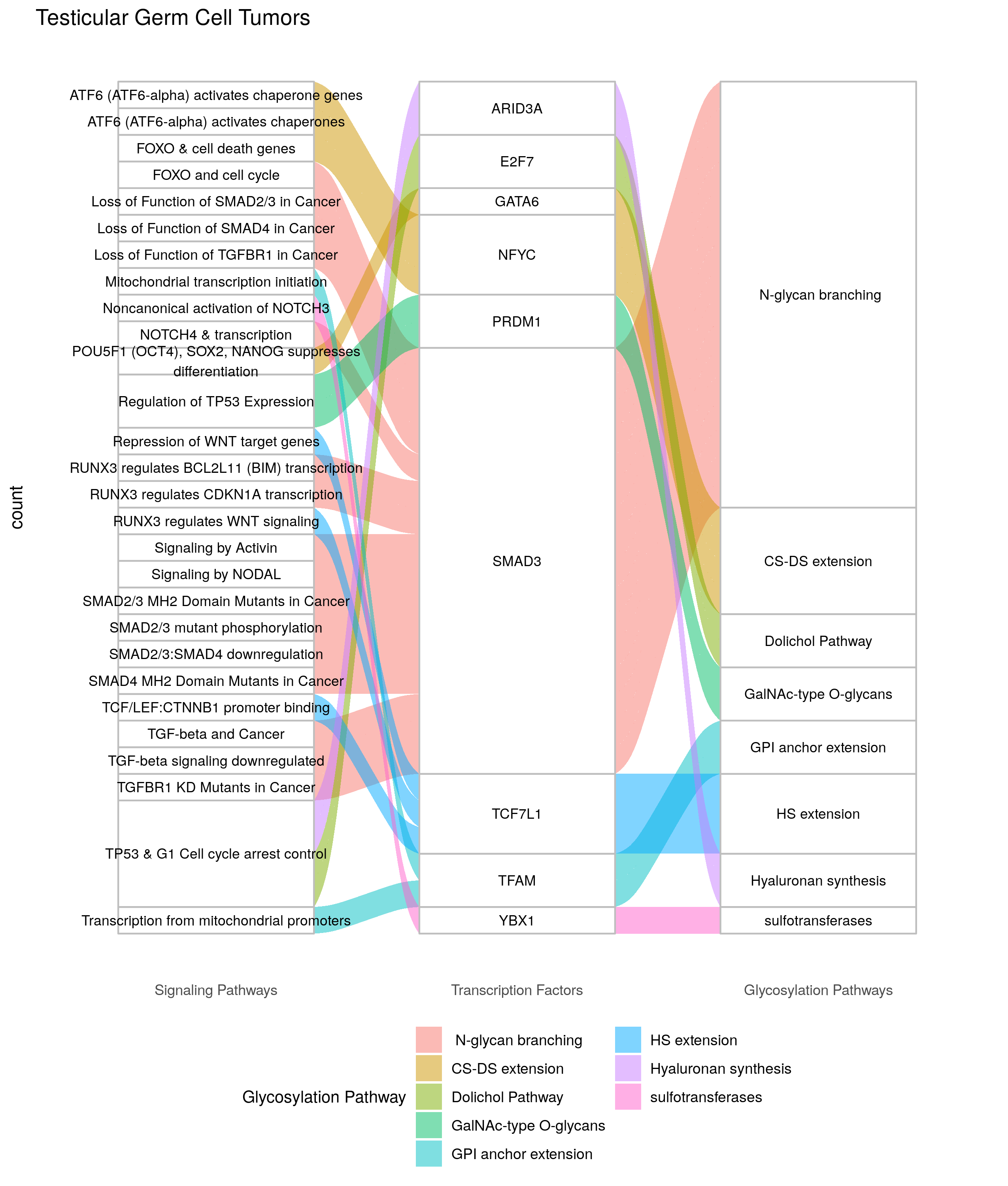

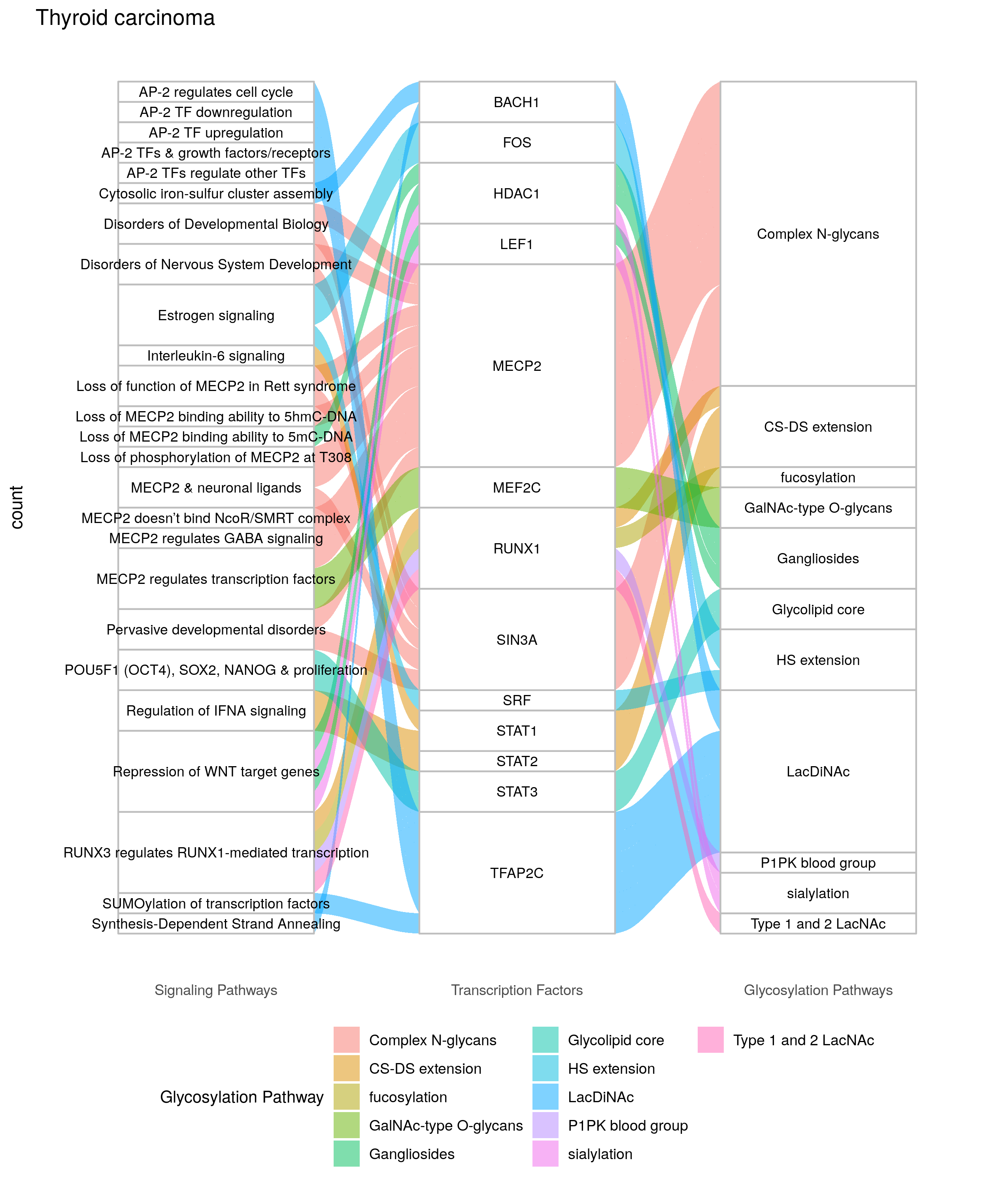

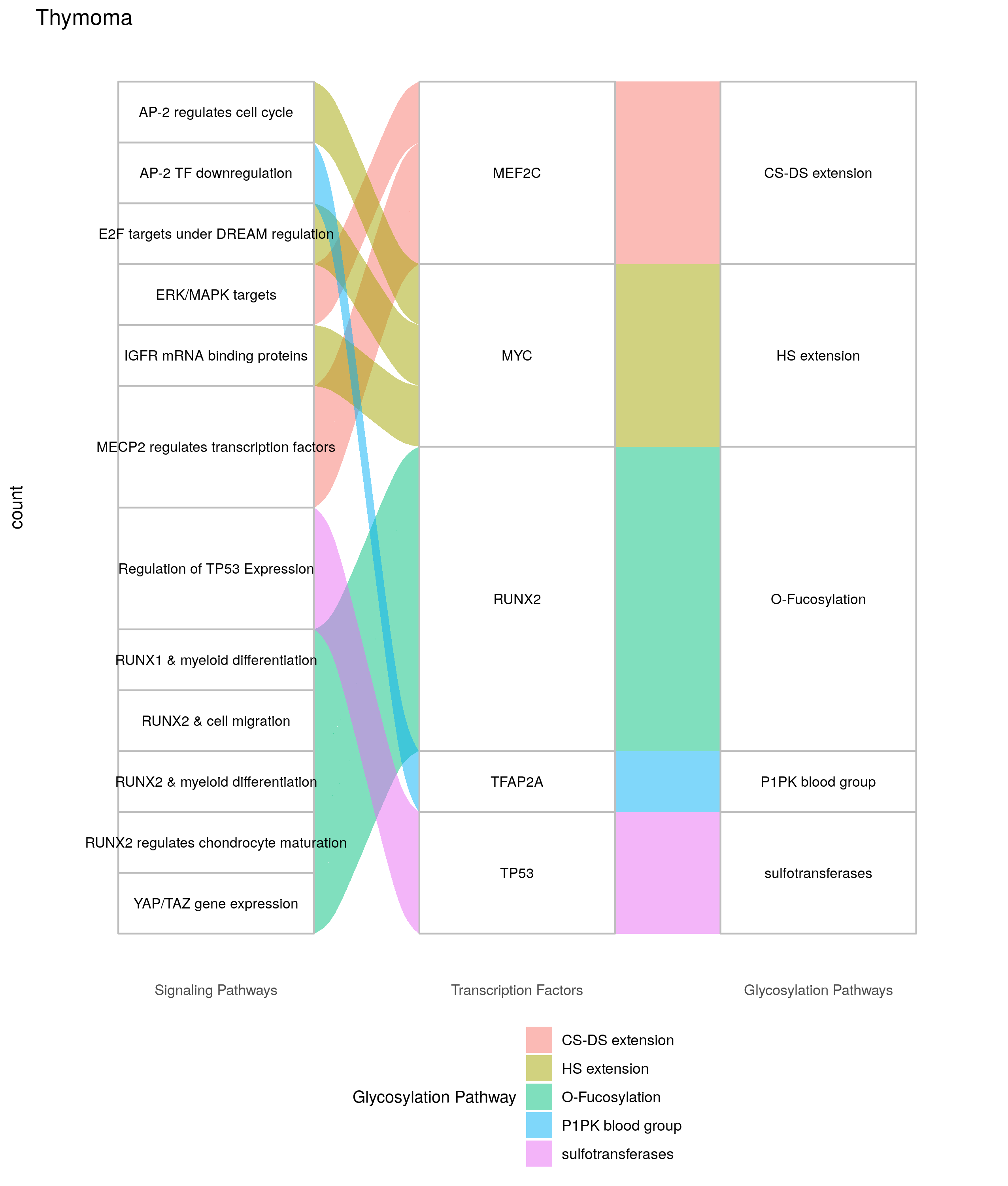

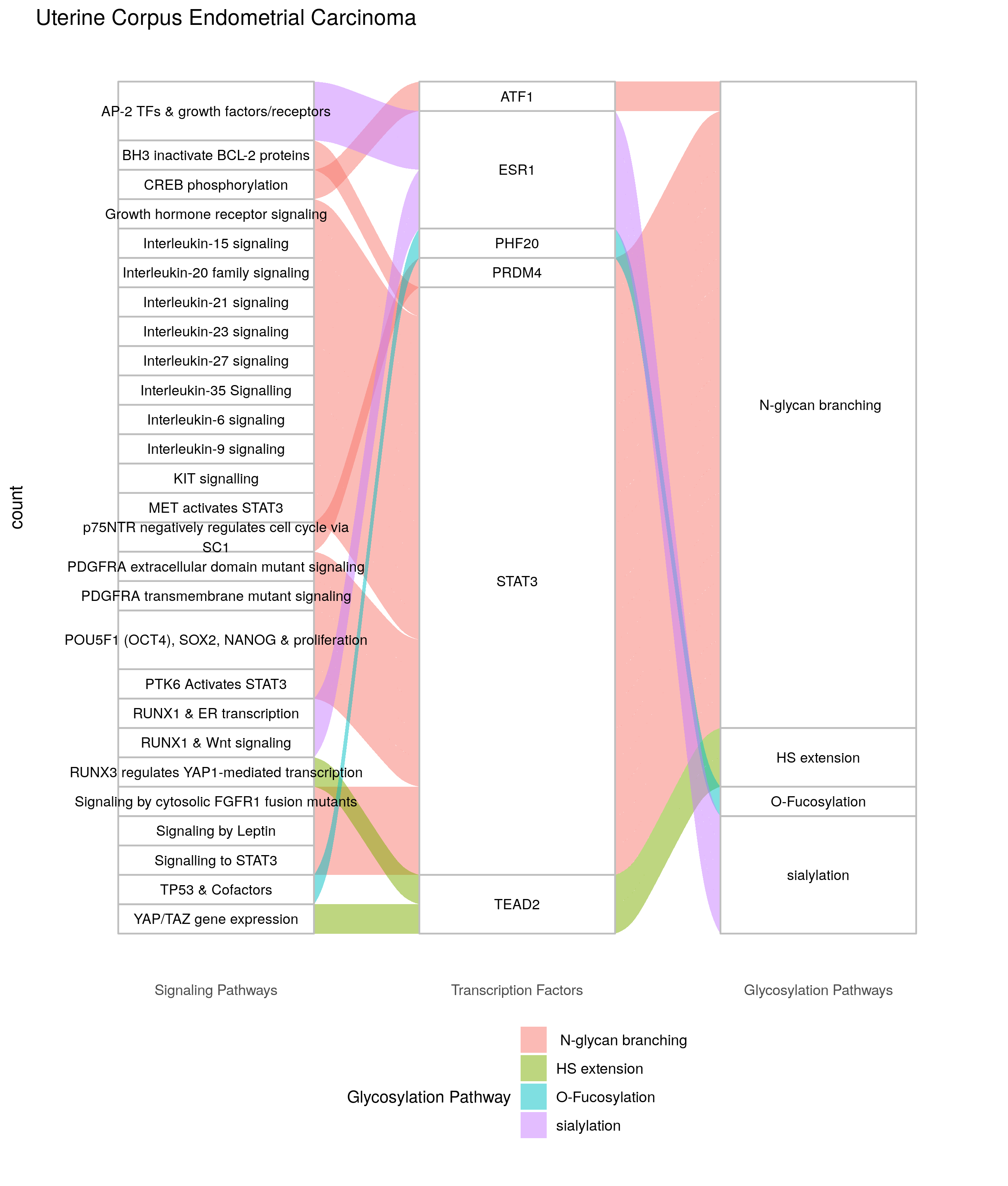

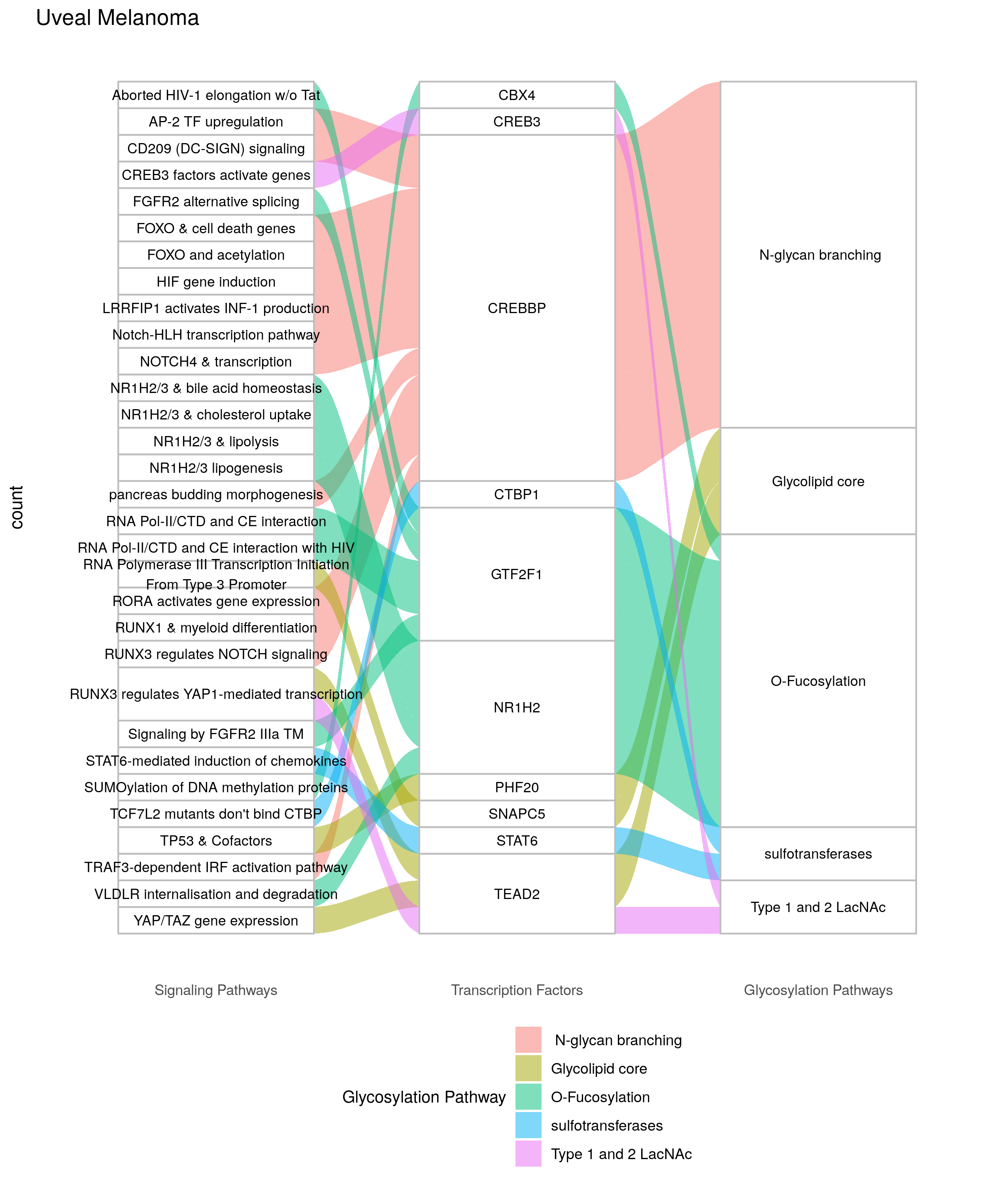
